# Kinetic logic of uridylation-mediated RNA decay

**DOI:** 10.64898/2026.03.27.714668

**Authors:** Annamaria Sgromo, Benjamin M. Jordan, Andrew Aarestad, David Mörsdorf, Franziska Boneberg, Martin Jinek, Thomas R. Burkard, Niko Popitsch, Stefan L. Ameres

**Affiliations:** Max Perutz Labs, Vienna BioCenter (VBC), Dr.-Bohr-Gasse 9, 1030, Vienna, Austria; University of Vienna, Max Perutz Labs, Dr.-Bohr-Gasse 9, 1030, Vienna, Austria; University of Konstanz, Systems Biology of Development, Universitätsstraße 10, 78464 Konstanz, Germany; Sense Ai, Inc., St. Paul, Minnesota, USA; University of Vienna, Department of Neurosciences and Developmental Biology, Djerassiplatz 1, 1030 Vienna, Austria; University of Zurich, Department of Biochemistry, Winterthurerstrasse 190, 8057 Zurich, Switzerland; Institute of Molecular Biotechnology, IMBA, Vienna BioCenter (VBC), Dr.-Bohr-Gasse 3, 1030 Vienna, Austria

**Keywords:** RNA surveillance, post-transcriptional regulation, RNA decay, enzyme kinetics

## Abstract

3′-terminal uridylation marks structured non-coding RNAs for cytoplasmic decay, yet how uridylation is quantitatively coupled to exonucleolytic degradation remains unclear. Here, we dissect the kinetic logic of uridylation-mediated RNA surveillance in *Drosophila melanogaster*. Using biochemical reconstitution together with high-throughput enzymology and quantitative modeling, we show that the terminal uridylyl transferase Tailor generates discrete oligo(U) intermediates through product-dependent kinetic tuning, while co-substrate promiscuity suppresses sustained processivity under physiological nucleotide conditions. Massively parallel binding and decay assays further reveal how the 3′-to-5′ exoribonuclease Dis3l2 selectively degrades Tailor-primed RNAs by integrating 3′-proximal uridine content and defined 3′-end accessibility—features encoded by short, kinetically tuned oligo(U) intermediates centered on four nucleotides—to enable productive threading of RNA substrates along an extended RNA-binding path to the catalytic site. Together, our findings establish a quantitative framework in which uridylation encodes decay competence through transient RNA 3′-end states that are matched to the mechanistic requirements for decay.

**Highlights:**

- The TUTase Tailor kinetically tunes uridylation to generate short, discrete oligo(U) intermediates
- Mixed nucleotide availability suppresses sustained processive uridylation
- Dis3l2 decodes 3′-proximal uridine content and end accessibility to commit RNAs to decay
- Short oligo(U) tails encode RNA decay competence through transient RNA 3′-end states

## Introduction

RNA quality control (QC) pathways safeguard gene expression by selectively identifying and eliminating aberrant transcripts through multiple, mechanistically distinct decay routes^1^. This task is particularly challenging for structured non-coding RNAs (ncRNAs), whose misprocessing can generate stable, highly folded molecules that are frequently resistant to exonucleolytic degradation. Effective surveillance therefore requires mechanisms that not only distinguish defective RNAs from functional species, but also actively modify RNA ends to render them competent for decay.

A conserved strategy to overcome the intrinsic decay resistance of structured RNAs relies on post-transcriptional 3′-end tailing, the non-templated addition of nucleotides by terminal nucleotidyl transferases, which functions as a regulatory signal for RNA processing and turnover^2-6^. The functional consequences of 3′ tailing are context dependent. In bacteria, adenylation can facilitate RNA decay by providing single-stranded extensions that promote exonucleolytic access, whereas in eukaryotes adenylation stimulates nuclear degradation of aberrant transcripts by the RNA exosome, while cytoplasmic polyadenylation stabilizes mRNAs and enhances translation^1,7,8^. In contrast, RNA uridylation, catalyzed by terminal uridylyl transferases (TUTases), has emerged as a prominent modification associated with cytoplasmic RNA surveillance across eukaryotes^6,9-13^. Notably, uridylation does not function as a uniform decay signal: in metazoans, mono-uridylation of specific precursor microRNAs enhances RNA maturation by restoring the optimal 3′-end geometry required for Dicer processing, whereas oligo-uridylation promotes degradation of defective or improperly processed RNAs^11,14-16^. Together, these observations indicate that the impact of uridylation on RNA fate depends on tail length, nucleotide composition, and RNA context, but how these parameters are regulated to selectively commit RNAs to decay remains poorly understood.

Uridylation-mediated RNA surveillance targets aberrant structured ncRNAs through the coordinated action of terminal uridylyl transferases and the 3′–5′ exoribonuclease Dis3l2. In *Drosophila melanogaster*, this theme is exemplified by the terminal RNA uridylation-mediated processing (TRUMP) pathway, in which the TUTase Tailor modifies structured RNAs and promotes their degradation by Dis3l2^17,18^. Tailor acts on a broad range of substrates, including splicing-derived mirtrons, mature microRNAs, and unprocessed RNA polymerase III transcripts^17,19,20^. Related uridylation-dependent surveillance of structured cytoplasmic ncRNAs by Dis3L2 has been described in mammals, where TUTases and the Perlman syndrome exonuclease Dis3L2 promote decay of vault RNAs, Y RNAs, snRNA precursors, and other aberrantly processed transcripts^21-23^. Despite this conservation, it remains unclear how uridylation is quantitatively tuned to generate RNA substrates that are productively engaged by Dis3l2, or why only a subset of uridylated RNAs are efficiently committed to decay.

Structural studies have provided important insight into the molecular architecture underlying uridylation-mediated RNA decay. Crystal structures of Tailor revealed an active-site configuration that favors uridylation initiation on specific 3′-terminal nucleotides, explaining aspects of substrate discrimination at the onset of tailing^24,25^. Complementary structural and cryo–electron microscopy analyses of mammalian and yeast Dis3L2 revealed an extended oligo(U)-binding trajectory that guides RNA from an open entry funnel to the catalytic site, spanning approximately 14 nucleotides and engaging uracil-specific contacts distributed across multiple binding tracts^26-29^. These studies further showed that Dis3L2 undergoes pronounced conformational rearrangements during RNA engagement, enabling helicase-independent degradation of structured substrates^27^. Together, these observations imply that uridylation must generate RNA 3′-ends with specific length, composition, and accessibility to support productive binding and threading through the catalytic channel; however, how such substrates are selectively generated and how Dis3L2 interrogates these features to commit RNAs to decay remain unresolved.

Across RNA tailing pathways, tail length has emerged as a critical determinant of RNA fate, reflecting the balance between distributive and processive polymerase activity. Trf4-Air2-Mtr4 polyadenylation (TRAMP) complex-mediated polyadenylation appears to operate distributively *in vitro*, whereas canonical poly(A) polymerases are processive, indicating how differences in catalytic mode can shape RNA processing outcomes^30^. Similarly, mammalian TUT4/7 enzymes undergo regulated shifts between mono-uridylation and oligo-uridylation in response to the RNA-binding protein Lin28, which promotes pre-let-7 uridylation and decay during development^14,16,31-33^. Together, these examples highlight kinetic tuning of terminal nucleotidyltransferase activity as a general mechanism for shaping RNA fate and suggest that uridylation-mediated RNA decay is governed not simply by the presence of a U-tail, but by a quantitative interplay between tail synthesis and downstream exoribonuclease recognition.

Here, we define how uridylation is quantitatively coupled to RNA decay. We show that the terminal uridylyl transferase Tailor generates discrete oligouridylated RNA intermediates through kinetically tuned tail synthesis rather than indiscriminate poly(U) extension. These intermediates encode decay competence by specifying uridine content, 3′-end accessibility, and the capacity of the RNA end to support productive threading into the exoribonuclease Dis3l2. Dis3l2, in turn, selectively commits RNAs to processive degradation by decoding these substrate features. Together, our findings establish a kinetic framework for uridylation-mediated RNA surveillance in which controlled tail synthesis creates RNA 3′-end states optimally matched to the mechanistic requirements of exoribonucleolytic decay.

## Results

### Tailor transitions from distributive to processive uridylation *in vitro*

To examine the enzymatic properties of Tailor, we expressed and purified recombinant (His)_6_-MBP-Tailor from *Sf9* insect cells (Figure 1A) and reconstituted RNA uridylation under physiologically relevant enzyme and UTP co-substrate concentrations (30 nM Tailor, 10 nM 5′-^32^P-labeled RNA substrate, 0.5 mM UTP; Figure S1A-C). Under these conditions, Tailor exhibited robust 3′ uridylation activity, resolved at single-nucleotide resolution by denaturing gel electrophoresis (Figure 1B). Early reaction time points revealed a pronounced accumulation of discrete oligo-uridylated intermediates predominantly containing three to five uridines, whereas extended incubations also resulted in the formation of long poly(U) tails rapidly exceeding 50 nucleotides. The coexistence of stable short intermediates and highly elongated products is reminiscent of kinetically separable catalytic modes corresponding to distributive and processive uridylation states, respectively.

**Figure 1.**
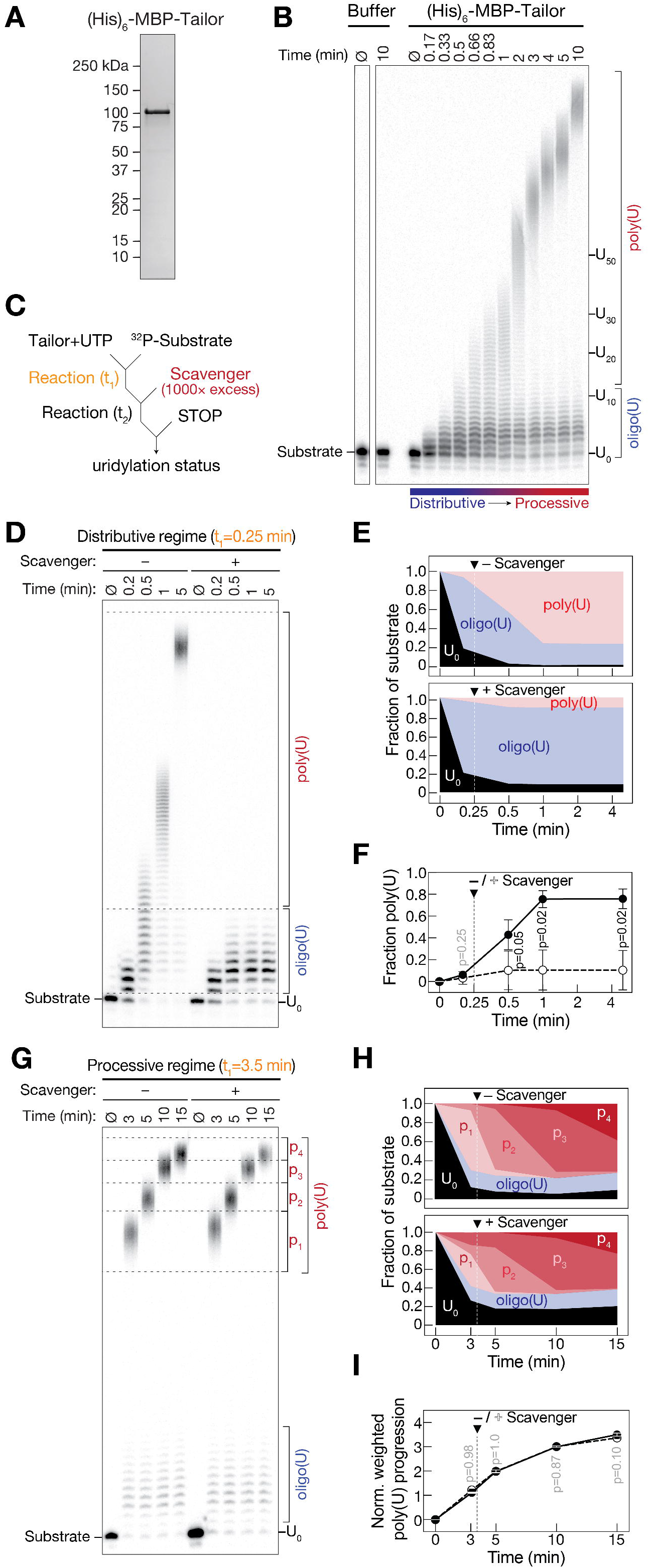
Tailor switches from distributive oligouridylation to processive polyuridylation. **(A)** Coomassie-stained SDS–PAGE of recombinantly purified (His)_6_–MBP–Tailor used for *in vitro* uridylation assays. **(B)** Time-course uridylation assay using a 5′-radiolabeled single-stranded RNA substrate incubated with recombinant (His)_6_–MBP–Tailor and UTP (0.5 mM). Reaction products were resolved by denaturing PAGE and visualized by phosphorimaging. Positions of untailed substrate (U_0_), oligouridylated RNA (oligo(U)), and polyuridylated RNA (poly(U)) are indicated. **(C)** Schematic of the scavenger assay used to assess uridylation processivity. After an initial reaction period (t_1_), a 1000-fold excess of unlabeled RNA scavenger is added before continuation of the reaction (t_2_). **(D)** Scavenger assay performed at an early time point (t_1_ = 0.25 min), corresponding to the distributive regime. Addition of scavenger prevents further tail elongation, indicating distributive oligouridylation. **(E)** Quantification of substrate, oligo(U), and poly(U) fractions over time for reactions shown in (D), with and without scavenger. **(F)** Comparison of poly(U) accumulation over time in the distributive regime (from D,E). Points represent mean ± SD. P-values were determined using Welch’s t-test with Benjamini–Hochberg correction. **(G)** Scavenger assay performed at a later time point (t_1_ = 3.5 min), corresponding to the processive regime. Poly(U) tails continue to elongate despite scavenger addition, indicating processive uridylation. **(H)** Quantification of substrate, oligo(U), and poly(U) fractions over time for reactions shown in (G), with and without scavenger. **(I)** Comparison of normalized poly(U) progression in the processive regime (from G,H). Mean ± SD is shown; statistical analysis as in (F).

To distinguish whether these kinetically separable uridylation regimes correspond to distributive or processive enzymatic states, we performed *in vitro* competition assays in which a 1000-fold excess of unlabeled scavenger RNA was added at defined time points and further extension of the radiolabeled substrate was monitored (Figure 1C). Addition of scavenger RNA early during the reaction – corresponding to the presumed distributive regime at t_1_ = 0.25 min – strongly suppressed continued uridine incorporation, resulting in the accumulation of short oligo(U) species and a pronounced reduction in poly(U) products (Figure 1D,E). Quantification revealed a decrease in the fraction of poly(U) RNA in the presence of scavenger relative to control reactions (Figure 1E). Consistently, quantitative analysis of uridylation patterns showed a significantly reduced progression into the poly-uridylated state upon scavenger addition (Figure 1F; p < 0.05, Benjamini-Hochberg multiple comparison corrected Welch’s one-sided t-test), indicating frequent enzyme dissociation during early uridylation cycles. In contrast, scavenger addition after accumulation of polyuridylated intermediates –corresponding to the processive regime at t_1_ = 3.5 min – did not impact further tail extension. Under these conditions, poly(U) species continued to elongate over time despite the presence of excess competitor RNA (Figure 1G). Quantitative analysis showed no difference in either the overall fraction of poly(U) RNA (Figure 1H) or the normalized weighted progression of poly(U) tail elongation across individual length classes (Figure 1I; p > 0.10, Benjamini-Hochberg multiple comparison corrected Welch’s one-sided t-test). Control reactions confirmed that Tailor did not change activity over time and that the scavenger efficiently outcompeted the radiolabeled RNA (Figure S1D,E). Our observations indicate that, once established, the late uridylation state is resistant to competition.

Together, these results indicate that progressive uridine addition during tail synthesis drives an intrinsic transition of Tailor from distributive oligouridylation to processive polyuridylation, reflecting a dynamic shift in enzymatic mode.

### Co-substrate promiscuity constrains processive RNA uridylation

Tailor and related TUTases have been reported to display relaxed nucleotide specificity *in vitro*, preferentially utilizing – beyond UTP – ATP over CTP as nucleotide donors, with minimal GTP-supported incorporation^17,20,34-36^. However, how such co-substrate promiscuity influences uridylation efficiency, tail composition, or reaction kinetics remains unclear. To address this, we performed *in vitro* tailing assays in the presence of all four nucleoside triphosphates (NTPs), supplied either at equal (0.5 mM each) or at physiological concentrations (0.5 mM UTP and GTP, 0.3 mM CTP, and 3 mM ATP;^37^) (Figure 2A). Compared with UTP-only reactions, both mixed-NTP conditions exhibited a delayed accumulation of long poly(U) products, with shorter oligo(U) species persisting for extended times before the appearance of extensive tail elongation (Figure 2A and Figure S2A). Quantification of poly(U) formation over time revealed a significantly slower increase in the poly(U) fraction under both mixed-NTP conditions relative to UTP-only reactions (Figure 2B; p < 0.03, Benjamini-Hochberg multiple comparison corrected Welch’s one-sided t-test). This kinetic delay was more pronounced under physiological NTP concentrations, indicating that the nucleotide environment quantitatively modulates the rate of Tailor-mediated uridylation and diminishes processivity.

**Figure 2.**
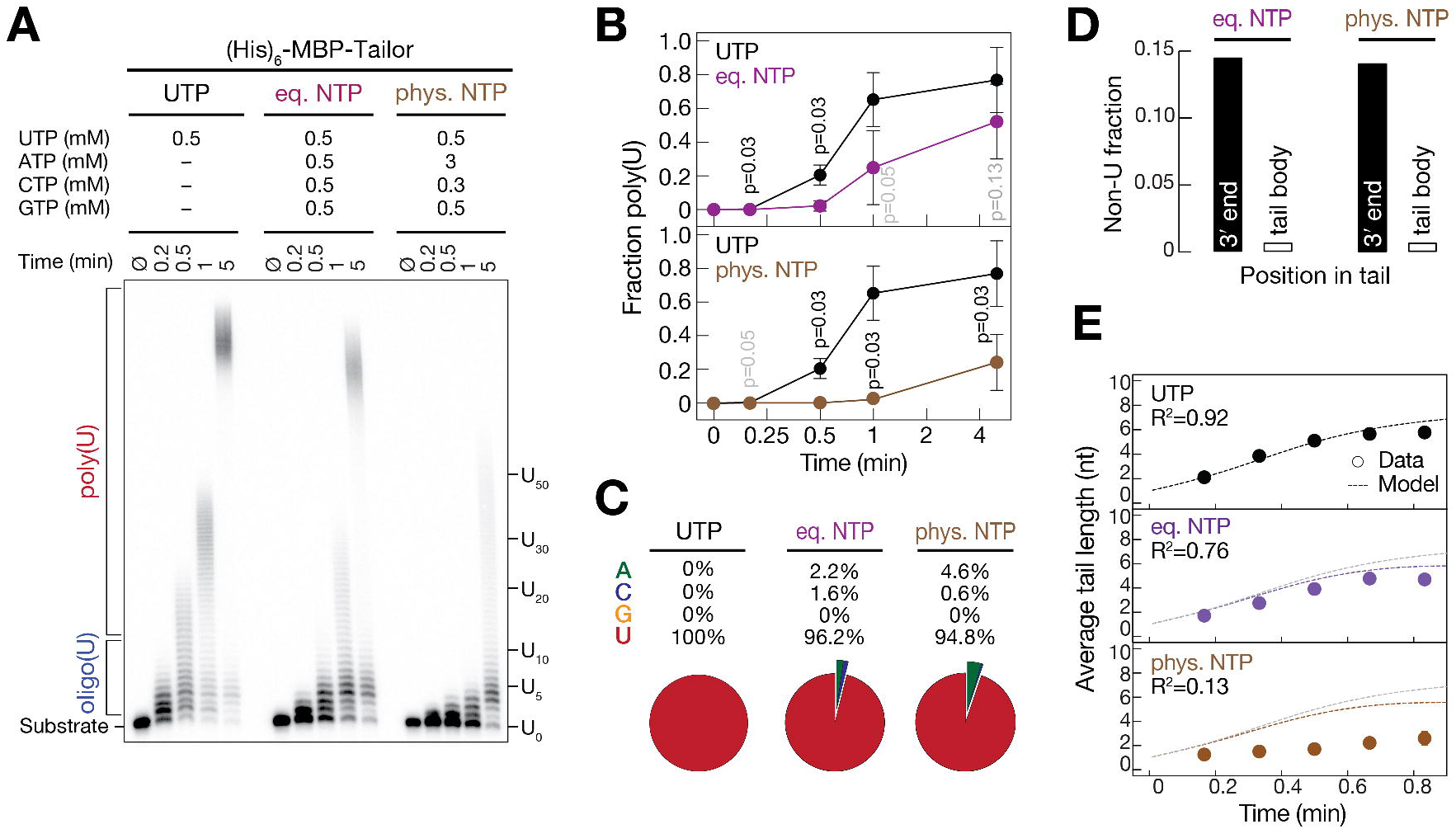
Co-substrate competition constrains Tailor processivity. **(A)** *In vitro* uridylation time course using recombinant (His)_6_–MBP–Tailor and a 5′-radiolabeled RNA substrate in the presence of UTP alone (0.5 mM), equal NTP concentrations (0.5 mM each), or physiological NTP concentrations (0.5 mM UTP and GTP, 0.3 mM CTP, 3 mM ATP). Reaction products were resolved by denaturing PAGE and visualized by phosphorimaging. **(B)** Quantification of poly(U) accumulation over time from assays in (A). Data represent mean ± SD from independent experiments; p-values from Welch’s t-test with Benjamini–Hochberg correction are indicated. **(C)** Nucleotide composition of uridylation products at 50 seconds, determined by high-throughput sequencing. Pie charts show the fraction of each incorporated nucleotide across all tail positions. **(D)** Fraction of non-uridine incorporation at the 3′ terminus versus the internal tail body for tails of length 2–10 nt under equal or physiological NTP conditions. **(E)** Average tail length over time derived from sequencing data (points) compared with simulations from a chain-termination model (lines). Model fits (R^2^) are shown for each nucleotide condition.

To directly examine nucleotide incorporation, we performed high-throughput sequencing of tailing products generated at early reaction timepoints (≤ 50 sec) in presence of equal or physiological NTP concentrations. As expected, uridine was the predominant incorporated nucleotide under all conditions (100% in UTP-only reactions, >96% with equal NTPs, and >94% under physiological NTP concentrations), confirming Tailor’s strong intrinsic preference for UTP (Figure 2C; Figure S2B-D). Position-resolved analysis of tail composition revealed that the infrequent non-uridine incorporation displayed a pronounced positional bias. Adenosine (up to 2.2% under equal NTPs and up to 4.6% under physiological NTP concentrations) and cytidine (up to 1.6% and 0.6 %, respectively) were detected predominantly at the 3′ termini of RNA tails, where non-U residues collectively accounted for up to 15% of 3′-terminal nucleotides (Figure 2C,D; Figure S2E-G). In contrast, non-uridine nucleotides were strongly depleted within the internal tail body (< 0.8%), indicating their incorporation is rarely followed by further nucleotide addition (Figure 2D; Figure S2H,I). This pronounced enrichment of non-uridine residues at tail 3′-ends together with the observed modulation of tail length progression (Figure 2A,B), suggests that sporadic non-U incorporation impedes continued tail extension by Tailor.

To evaluate whether chain termination by non-U incorporation alone could account for the observed reduction in tail length, we modeled uridylation kinetics with and without chain-terminating events and compared predicted average tail lengths to those measured by high-throughput sequencing (Figure 2E). At the experimentally observed low frequency of non-U incorporation, the model predicted a reduction in average tail length that was largely consistent with the data obtained under equal NTP conditions (R^2^ = 0.76). In contrast, under physiological NTP concentrations, the experimentally observed reduction in tail length was substantially greater than predicted by sporadic chain termination alone, resulting in a poor model fit (R^2^ = 0.13), consistent with additional inhibition arising from non-productive co-substrate competition by excess ATP.

Together, these results indicate that Tailor-dependent uridylation is shaped by both non-uridine incorporation that limits tail extension and non-productive co-substrate binding that slows uridylation kinetics.

### High-throughput analysis of stepwise Tailor uridylation kinetics

Bulk uridylation assays revealed a pronounced accumulation of oligo-uridylated intermediates containing three to five uridines (Figure 1B and 2A), suggesting that Tailor extends RNA tails through discrete kinetic steps rather than uniform elongation. To resolve these stepwise kinetics at single-nucleotide resolution, we analyzed uridylation of a 39-nt single-stranded RNA substrate bearing six randomized nucleotides at its 3′-end (6N RNA; 4,096 unique substrates; Figure 3A). We sampled tailing reactions at defined time points, resolved products by denaturing PAGE, and in parallel subjected each time point to high-throughput sequencing (HTS) (Figure 3B).

**Figure 3.**
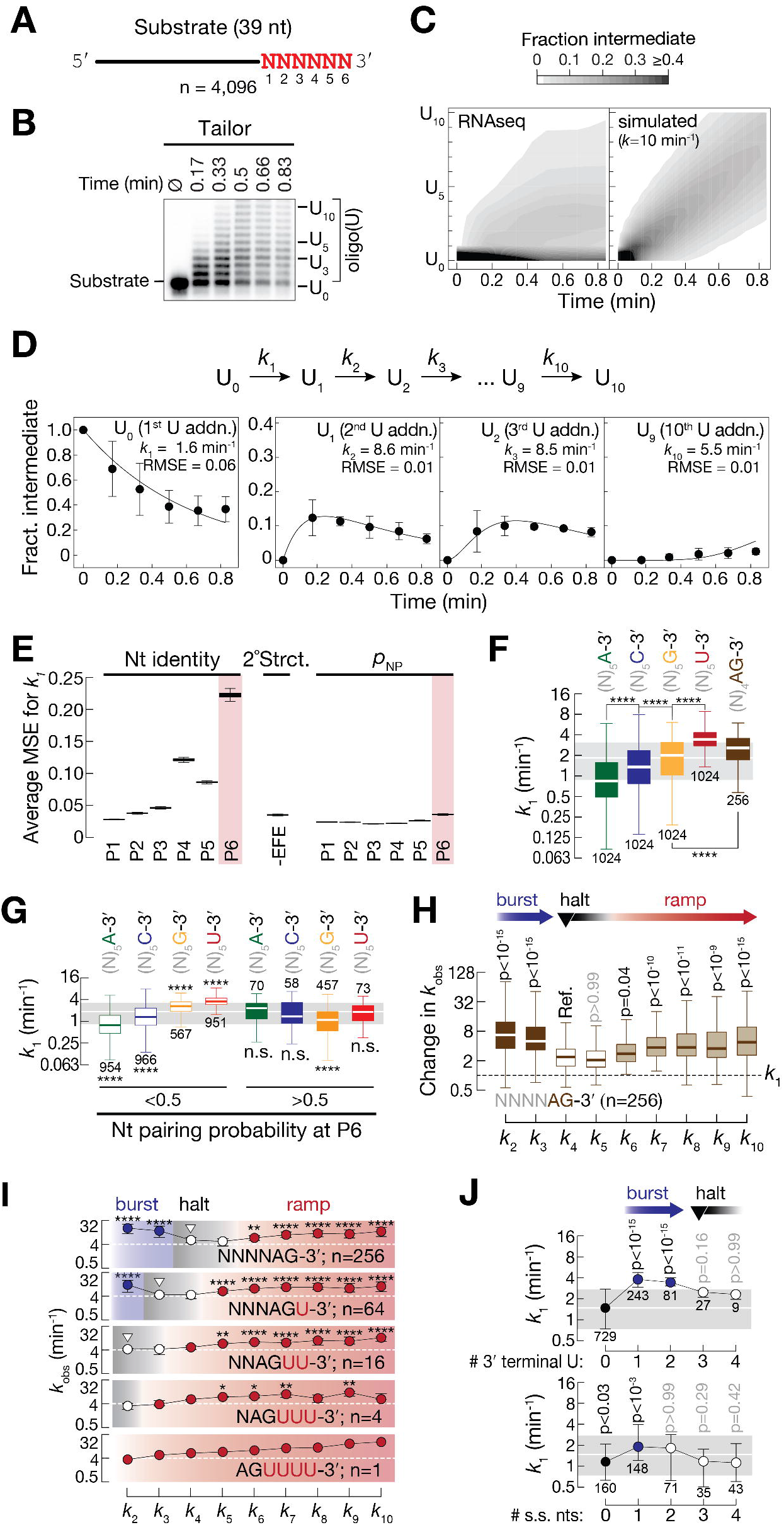
Tailor senses emerging U-tail length to kinetically tune oligouridylation. **(A)** Sequence of the single-stranded RNA substrate containing six randomized nucleotides at the 3′-end (6N RNA) used for high-throughput tailing assays. **(B)** Representative denaturing PAGE showing time-resolved oligouridylation (U_0_–U_10_) of the 5′-radiolabeled RNA substrate in (A) by Tailor. At each time point, RNA was recovered for high-throughput sequencing to quantify uridylation intermediates. **(C)** Contour plots showing the temporal distribution of uridylated intermediates. Left, experimentally measured fractions of summed normalized substrates derived from sequencing. Right, simulated uridylation assuming equal rate constants for all steps (*k*_1-10_ = 10 min^−1^). **(D)** Stepwise kinetic model of irreversible oligouridylation (top). Time courses for representative intermediates (U0, U1, U2, U9) averaged across all 4,096 substrates are shown (mean ± SD; n = 3 independent experiments). Solid lines indicate model fits; fitted rate constants (*k*_obs_) and root-mean-square errors (RMSE) are indicated. See Materials and Methods for the fitting equations and details of the analysis. **(E)** Permutation feature importance analysis for the prediction of the first uridylation rate (ln(1 + k_1_)). Input features include nucleotide identity (b1), base-pairing probability at each 3′ position (pN1), and ensemble free energy (−EFE). Mean squared error (MSE) increase upon feature permutation is shown (25 permutations per feature). **(F)** Distribution of *k*_1_ values for substrates grouped by 3′-terminal nucleotide identity. Box plots show median and interquartile range; numbers indicate substrates per group. Adjusted p-values were calculated using Kruskal–Wallis test with multiple-comparison correction relative to all substrates; ****, p < 10^-4^. **(G)** *k*_1_ values for substrates grouped by terminal nucleotide identity and stratified by predicted base-pairing probability at position P6 (>0.5 or <0.5). Box plots show median and interquartile range; numbers indicate substrates per group. Adjusted p-values were calculated using Kruskal–Wallis test with multiple-comparison correction relative to all substrates; ****, p<10^-4^; n.s., p > 0.05. **(H)** Relative uridylation rates for substrates terminating in −AG, normalized to *k*_1_. Adjusted p-values were calculated using Kruskal–Wallis test with multiple-comparison correction relative to the reference step (*k*_4_). **(I)** Uridylation rates (*k*_2_–*k*_10_) for substrates ending in −AG (top) or −AGU, −AGUU, −AGUUU, or −AGUUUU (bottom), mimicking progressively uridylated intermediates. Dashed lines indicate the mean rate for the −AG reference; white triangles denote the halt condition. Adjusted p-values were calculated using Kruskal–Wallis test; *, p<0.05; **, p<0.01; ***, p<10^-3^; **** p<10^-4^. **(J)** Uridylation rate *k*_1_ as a function of terminal uridine number (top) or predicted number of unpaired 3′-terminal nucleotides (bottom). Box plots show median and interquartile range; adjusted p-values were calculated using Kruskal–Wallis test relative to substrates lacking terminal uridines. Numbers under each error bar represent the number of substrates in each group.

Consistent with qualitative analyses, HTS revealed a characteristic temporal progression of uridylation intermediates, with discrete species accumulating sequentially over time rather than a continuous redistribution of tail lengths (Figure 3C, left). Intermediates carrying three to five uridines reached maximal abundance at intermediate time points before declining as longer tails accumulated. Gel-based quantification independently reproduced this enrichment of short intermediates (Figure S3A), confirming close agreement between sequencing- and PAGE-based measurements. This observation contrasts with simulations assuming constant uridylation rates that predicted a smooth, monotonic shift in tail-length distributions without intermediate accumulation (Figure 3C, right). Together, these observations show that Tailor-mediated oligouridylation proceeds through discrete, kinetically resolvable intermediates rather than uniform tail extension.

To quantitatively describe these dynamics, we modeled oligouridylation as a series of irreversible pseudo–first-order reactions and fitted step-specific rate constants (*k*_1_–*k*_10_) directly to the HTS time-course data (Figure 3D; see Materials and Methods). The fitted model closely reproduced the experimentally measured intermediate abundances across all time points and substrates, yielding low residual errors and stable parameter estimates (RMSE < 0.06; Figure 3D and Figure S3B). These results show that the kinetic framework faithfully captures Tailor-mediated oligouridylation and supports robust estimation of step-resolved uridylation rates.

### 3′-end sequence and local structure govern initial substrate engagement by Tailor

We next exploited the sequence diversity of the 6N substrate pool to identify RNA features that govern Tailor activity. Using HTS-derived rate constants, we trained machine-learning models incorporating substrate features, including nucleotide identity, predicted RNA secondary structure (effective free energy, EFE), and base-pairing probability (*p*_*NP*_). The models achieved high predictive performance, with strong agreement between predicted and observed values (R^2^ = 0.90; Pearson r = 0.96; Spearman ρ = 0.96; Figure S3D) and low training and validation losses (Figure S3E). Across all models, the identity of the terminal nucleotide (position P6; Figure 3A) emerged as the dominant determinant of the first uridylation step (*k*_1_), which we used as a proxy for initial substrate engagement (Figure 3E and Figure S3C–E).

Consistent with these predictions, analysis of the first uridylation step (*k*_1_) showed that Tailor uridylated RNAs terminating in 3′-U and 3′-G significantly faster than substrates ending in 3′-A or 3′-C, revealing a clear ordering of substrate engagement among terminal nucleotides (Figure 3F; p < 10^−4^, Kruskal–Wallis test with multiple-comparison correction). Substrates containing a 3′-AG motif were enriched among RNAs with high *k*_1_ values, consistent with previous *in vitro* and *in vivo* observations identifying splice acceptor sites as frequent Tailor targets^17,19,20,24,25^. Sequence-logo analysis of the top and bottom 10% of substrates ranked by *k*_1_ further showed enrichment of 3′-terminal uridine among the most efficiently tailed RNAs and enrichment of 3′-terminal adenosine among the least efficiently tailed RNAs (Figure S3F).

To assess how RNA structure influences the first uridylation step (*k*_1_), we first examined correlations across all substrates. *k*_1_ showed only weak but statistically significant negative correlations with both overall RNA secondary structure (negative effective free energy, -EFE; Spearman r = −0.11, p < 10^−4^) and with base-pairing probability at the 3′-terminal nucleotide (weighted Spearman r = −0.07; Figure S3G,H), indicating that neither metric alone strongly predicts initial Tailor engagement. We therefore analyzed pairing probability at the terminal position (P6) in a nucleotide-resolved manner (Figure S3I). This analysis revealed opposing terminal-nucleotide– specific responses: increased P6 pairing correlated negatively with *k*_1_ for substrates ending in 3′-G (r = −0.57) and 3′-U (r = −0.17), and positively for substrates ending in 3′-A (r = +0.57) and 3′-C (r = +0.15). Because of these opposing effects, grouping substrates by paired versus unpaired terminal nucleotides showed convergence of *k*_1_ values across all terminal nucleotide classes when the 3′ terminus was paired (Figure 3G), demonstrating that 3′-end pairing neutralizes terminal-nucleotide– specific effects. Stratification by overall RNA secondary structure indicated that global folding contributes little explanatory power beyond its influence on terminal accessibility (Figure S3J); and position-resolved analysis confirmed that correlations between pairing probability and *k*_1_ are confined to the terminal nucleotide itself, with minimal contributions from upstream positions (P1– P5) (Figure S3K,L).

### Emerging U-tails shape Tailor uridylation kinetics

Building on the step-resolved kinetic framework described above, we analyzed how uridylation rates evolve after initial substrate engagement by examining the rate constants governing successive uridine additions (*k*_2_–*k*_10_). For substrates terminating in a 3′-AG motif—a class that is efficiently engaged by Tailor and representative of physiological 3′-ends—step-specific rate constants followed a reproducible, non-monotonic pattern (Figure 3H), with rapid uridine addition during the early steps (*k*_2_-*k*_3_; ‘burst’), pronounced attenuation at intermediate steps (*k*_4_-*k*_5_; ‘halt’), and stepwise re-acceleration at later steps (*k*_6_-*k*_10_; ‘ramp’). Early, intermediate, and late rate constants differed significantly (adjusted p < 0.04, Kruskal–Wallis test with Dunn’s multiple comparison correction), indicating that Tailor-mediated uridylation does not proceed with constant rates. At later stages of tail extension, step-specific rate constants became more similar, suggesting reduced step dependence of uridylation kinetics as tails lengthen and successive uridine additions become more uniform.

To test whether the observed kinetics reflect a general response to progressive tail growth rather than a property of a specific 3′-end sequence, we analyzed substrates engineered to carry defined numbers of pre-existing terminal uridines. These substrates reproduced the same burst– halt–ramp pattern, with the position of the kinetic slowdown shifting according to terminal uridine content (Figure 3I). Extending this analysis beyond 3′-AG substrates, revealed a stereotypic modulation of step-specific rate constants across diverse sequence contexts (Figure S3M). These results define RNA substrate U-tail length as a general determinant of uridylation kinetics.

To distinguish whether stepwise kinetic modulation reflects sensing of the emerging U-tail or progressive exposure of single-stranded RNA at the 3′-end, we analyzed uridylation rates (*k*_1_) as a function of terminal uridine content and accessibility. Grouping substrates by pre-existing 3′-terminal uridines revealed significant differences in uridylation rate constants across sequence contexts (Figure 3J; p < 10^−15^, Kruskal–Wallis test with Dunn’s multiple comparison correction), capturing the characteristic burst of the first two uridine additions after initiation. In contrast, 3′-terminal overhangs exerted a limited effect: substrates with a single unpaired nucleotide at the 3′-end showed enhanced initial uridylation rates (p < 10^−3^, Kruskal–Wallis test with Dunn’s multiple comparison correction), whereas fully paired ends or longer overhangs did not support a significant further rate increase (Figure 3J). These observations indicate that nucleotide identity of a growing U-tail is the dominant determinant of kinetic modulation during oligouridylation.

Together, these results indicate that Tailor senses the growing U-tail and kinetically shapes uridylation in a ruler-like fashion, resulting in preferential accumulation of short, defined oligo(U) tails with an average length of ∼4 nts.

### Quantitative framework for Dis3l2 substrate engagement and decay

To determine how Tailor-generated oligo(U) intermediates enter the decay pathway, we examined RNA substrate recognition and degradation by Dis3l2 using the 39-nt RNA substrate bearing six randomized nucleotides at its 3′-end (6N RNA; 4,096 unique sequences) employed for Tailor uridylation assays (Figure 4A). We quantified RNA decay kinetics as well as binding affinities to enable direct integration of substrate engagement and decay.

**Figure 4.**
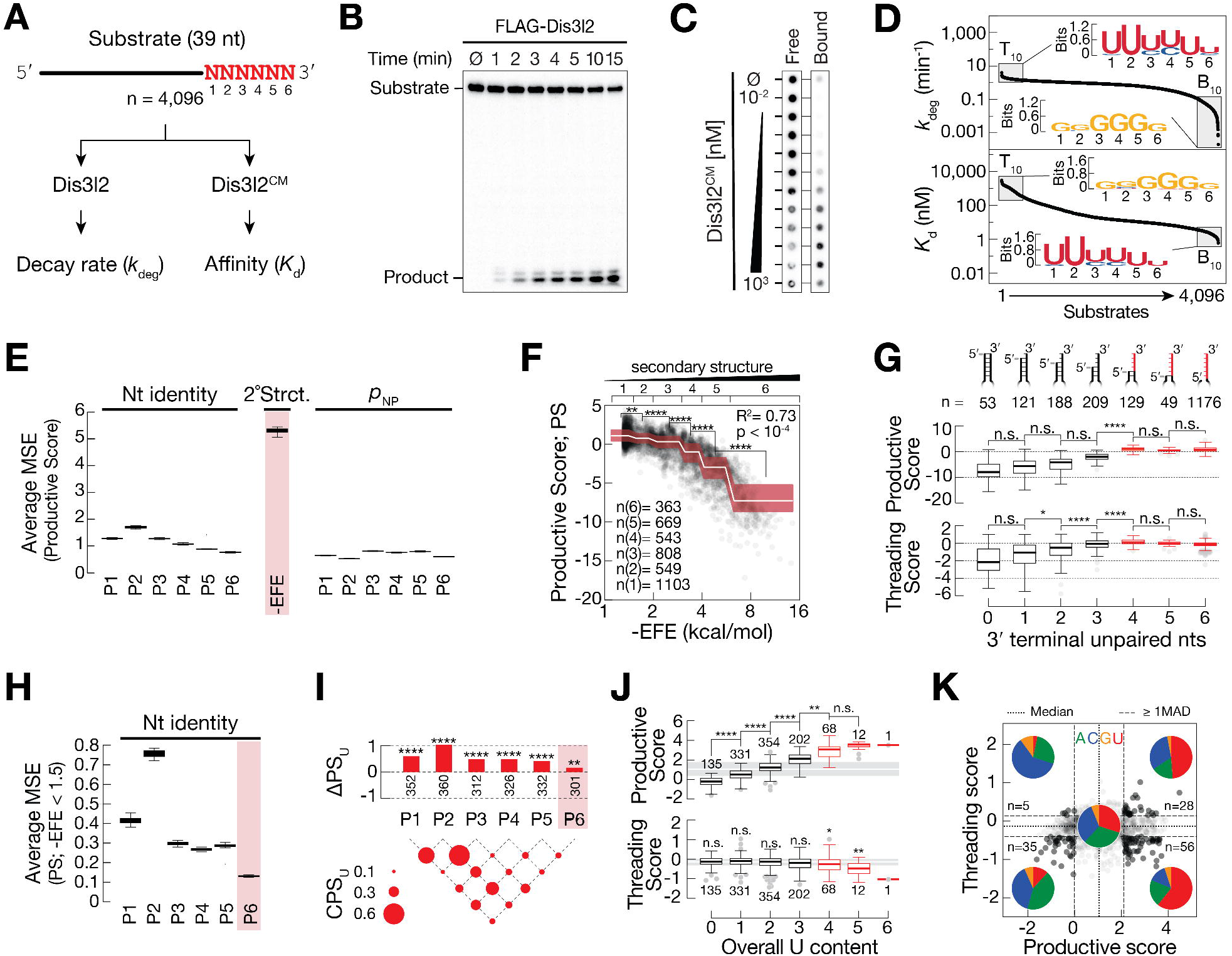
Dis3l2 integrates 3′-proximal uridine content and end accessibility to enable productive threading and decay. **(A)** Experimental overview of massively parallel degradation assays using wild-type Dis3l2 and RNA Bind-n-Seq assays using catalytically inactive Dis3l2^CM^ to derive decay rates (*k*_deg_) and dissociation constants (*K*_d_), respectively, for a 6N substrate pool (4,096 species). **(B)** Representative *in vitro* degradation assay of a radiolabeled 6N substrate by FLAG-Dis3l2 resolved by denaturing PAGE. **(C)** RNA Bind-n-Seq assay showing concentration-dependent binding of Dis3l2^CM^ to the 6N substrate pool. **(D)** Global distributions of *k*_deg_ (top) and *K*_d_ (bottom) values derived from sequencing-based fitting. Sequence logos for the top and bottom 10% of substrates are shown. **(E)** Relative permutation feature importance for prediction of productive score (PS), highlighting contributions of nucleotide identity, secondary structure, and ensemble folding energy (−EFE). **(F)** Relationship between RNA secondary structure (−EFE) and productive score (PS). Substrates were grouped into approximately evenly sized bins based on −EFE, and differences in PS between bins were statistically assessed using a Kruskal–Wallis test. PS correlates strongly with −EFE (R^2^ = 0.73; p < 10^−4^). Pairwise statistical significance between secondary structure bins is indicated; **** p < 10^−4^. **(G)** Tukey boxplots show Productive Score (top) and Threading Score (bottom) as a function of the number of unpaired 3′-terminal nucleotides. *P*-values were determined by Kruskal-Wallis test; n.s, p > 0.05; *, p < 0.05; ****, p<10^-4^. **(H)** Feature importance analysis (see panel E) restricted to single-stranded substrates. **(I)** Position-specific (top) and pairwise contributions (bottom) of uridines to productive score (PS) in unstructured RNAs. ΔPS indicates median shifts, and Cooperativity of Productive Score (CPS) quantify synergistic effects of uridines at the indicated positions (P1-P6) and combinations. Significance was determined by two-sided Mann–Whitney U test; **, p < 0.01; ****, p< 10^-4^. **(J)** Productive score (PS) and threading score (TS) for single-stranded substrates grouped by total uridine content at the 3′-end. Significance was determined by Kruskal–Wallis with Dunn’s correction. n.s., p > 0.05; *, p < 0.05; **, p < 0.01; ****, p < 10^-4^. **(K)** Productive score versus threading score highlighting outlier substrates (±1 median absolute deviation; MAD). Pie charts indicate nucleotide composition across the four quadrants.

Decay rate constants (*k*_deg_) were determined by high-throughput sequencing of *in vitro* decay reactions reconstituted with immunopurified FLAG-Dis3l2 (Figure 4B,D). Replicate experiments yielded highly reproducible decay profiles and robust kinetic fits across the dataset for 4,034 substrates (Figure S4A,B), with *k*_deg_ values spanning more than four orders of magnitude around a median decay rate of 0.53 min^−1^ (Figure 4D and Figure S4A).

In parallel, we measured RNA binding affinities (*K*_d_) using a recombinant, catalytically inactive Dis3l2 mutant (D568A, Δ51–200; hereafter Dis3l2^CM^), which retains RNA-binding capacity but lacks nuclease activity (Figure 4C,D). Sequencing-based quantification of protein-bound RNA across increasing Dis3l2^CM^ concentrations, combined with spike-in normalization, enabled reproducible affinity estimations for 4,096 substrates (Figure S4C-F). Binding affinities spanned more than four orders of magnitude around a median *K*_d_ of 26.5 nM (Figure 4D and S4E). Comparison of the most rapidly degraded and most tightly bound substrates revealed strong enrichment for uridine-rich sequences, whereas poorly bound and slowly degraded RNAs were depleted of uridines (Figure 4D), confirming prior biochemical and *in vivo* studies of Dis3L2^17,21,22,26,27,29,38-41^.

To integrate binding affinity and decay rate constants, we standardized both parameters using robust Z-scores. Standardized binding and decay measurements showed an overall positive correlation across substrates (R^2^=0.83, p<10^-4^; Figure S4G), indicating that stronger binding generally supports faster degradation while remaining insufficient to fully predict substrate-specific decay efficiency.

For further analyses, we defined a productive score (PS = Z[*k*_deg_] – Z[*K*_d_]) to identify substrates that combine strong binding with efficient degradation. Furthermore, we defined a threading score (TS = Z[*k*_deg_] + Z[*K*_d_]), which reports decay efficiency relative to binding strength, because efficient decay requires productive RNA translocation along a channel to the active site. Together, these composite metrics decouple RNA binding, threading, and catalytic efficiency, providing a quantitative framework for dissecting Dis3l2 substrate selection.

### 3′-end accessibility defines productive substrate binding and threading by Dis3l2

To determine how RNA substrate features shape productive Dis3l2 engagement beyond binding affinity alone, we analyzed determinants of both productive score (PS) and threading score (TS) across the full substrate pool. Machine-learning models trained on RNA sequence, predicted secondary structure, and local base-pairing probabilities identified overall RNA secondary structure as the dominant predictor of PS (Figure 4E; Figure S4H–J). Consistent with this result, PS inversely correlated with predicted folding stability, with less structured RNAs exhibiting higher productive engagement (Figure 4F; R^2^ = 0.73, p < 10^−4^). These analyses confirm that RNA accessibility is an important prerequisite for efficiently coupling Dis3l2 binding to downstream decay^17,26,27^.

To refine how local 3′-end accessibility contributes to Dis3l2 activity, we analyzed productive score (PS) and threading score (TS) as a function of the number of unpaired nucleotides at the 3′ terminus (Figure 4G). PS remained low for substrates with zero to three unpaired terminal nucleotides and increased most strongly and significantly between three and four unpaired nucleotides (p < 10^−4^; Kruskal–Wallis test with Dunn’s multiple-comparison correction), after which it plateaued. TS showed a significant increase as the number of unpaired 3′-terminal nucleotides increased but similarly reached maximal values at four unpaired nucleotides (p < 10^−4^; Kruskal– Wallis test with Dunn’s multiple-comparison correction) and did not further increase with additional unpairing. Thus, within the sequence space sampled here, both productive engagement and efficient threading converged once approximately four 3′-terminal nucleotides were accessible, while additional unpairing within this region did not measurably enhance Dis3l2 activity.

### A combinatorial 3′-proximal uridine logic governs Dis3l2 substrate engagement

To resolve sequence-dependent effects on Dis3l2 independent of global folding constraints, we restricted subsequent analyses to minimally structured RNAs (−EFE ≤ 1.5; n=1,103). Decay rate constants and binding affinities within the same substrate pool showed again a strong positive correlation (R^2^ = 0.71, p < 10^−4^; Figure S4K), yet substantial dispersion around this relationship remained, indicating that binding strength alone does not fully account for substrate-specific differences in decay efficiency and motivating a detailed analysis of position-dependent sequence effects. As expected, nucleotide identity exerted strong position-specific effects on Dis3l2 activity: feature-importance analysis identified uridines at 3′-proximal positions as the dominant contributors to high productive score (PS), with the strongest effect at position P2 and P1, whereas the 3′-terminal nucleotides (P6) contributed minimally (Figure 4H and Figure S4L–O).

To quantify positional nucleotide contributions systematically, we calculated a positional effect score (ΔPS) for each nucleotide at positions P1–P6. Uridine exhibited significantly positive ΔPS values at internal positions (p<10^-4^; Mann-Whitney U test), with the strongest contributions at P1 and P2, whereas the terminal position (P6) contributed minimally (Figure 4I and Figure S4P). In contrast, adenosine, cytidine, and guanosine showed predominantly negative ΔPS values across internal positions, indicating reduced productive engagement when non-uridine nucleotides occupy the 3′-proximal region (Figure S4P and Q). Importantly, none of the nucleotides—including uridine— showed a strong ΔPS contribution at the 3′-terminal position (P6), indicating that productive Dis3l2 engagement is largely insensitive to terminal nucleotide identity and instead depends on 3′-proximal sequence composition (Figure 4I and Figure S4Q).

We next assessed whether uridines act independently or cooperatively by calculating a uridine cooperativity score (CPS), which measures deviations from expected additive effects between pairs of uridines (Figure 4I, bottom). CPS analysis revealed statistically significant cooperativity across multiple position pairs, with the strongest cooperative contributions arising from 3′-proximal combinations, particularly P1–P2 and pairs involving P3 (p < 0.05; Mann–Whitney U test; Figure 4I). In contrast, cooperativity involving the terminal position (P6) was consistently weakest.

Taken together, Dis3l2 engagement follows a positional substrate-recognition logic in which uridines broadly promote decay but act most effectively when positioned proximally and in specific combinations, while the 3′-terminal nucleotide identity contributes minimally to productive engagement.

### Balancing uridine-dependent affinity and threading during Dis3l2-mediated decay

We next asked how uridine content distributes substrates across regimes dominated by productive engagement (PS) versus threading (TS). Increasing uridine number within the randomized region increased PS significantly up to four uridines (*p* < 0.01; Kruskal–Wallis with Dunn’s correction; Figure 4J, top) and then plateaued, whereas TS remained high through four uridines but declined significantly with further uridine enrichment (*p* < 10^-4^; Kruskal–Wallis with Dunn’s correction; Figure 4J, bottom). Thus, substrates with four uridines maximize productive engagement without incurring a threading penalty, while five or more uridines shift substrates toward a threading-limited regime.

To relate differences in productive engagement and threading to RNA sequence features, we mapped substrates according to their relative productive and threading scores and classified quadrant-enriched sequence classes using median absolute deviation–based outliers (Figure 4K and Figure S4R–T). Nucleotide-enrichment analysis (Figure 4K and Figure S4T) revealed distinct compositional biases across classes. Cytidine was enriched among substrates with efficient threading but reduced productive engagement (high TS, low PS; p < 0.01; two-sided binomial test), whereas adenosine was enriched among substrates with low scores in both dimensions (low PS, low TS; p < 0.01; two-sided binomial test), indicating poor performance in both binding-associated engagement and translocation. In contrast, uridine was strongly enriched among substrates with high productive engagement (p < 10^-4^; two-sided binomial test; Figure S4T), with sequence-logo analysis revealing that substrates in the high-PS/high-TS class preferentially contained short runs of consecutive uridines—most frequently four—within the 3′-proximal region (Figure 4K and Figure S4R-T). Substrates with longer uridine tracts (≥5 uridines) shifted predominantly into the high-PS/low-TS class, consistent with reduced threading efficiency beyond four uridines (Figure 4J and Figure S4R,S). Across all comparisons, the 3′-terminal nucleotide did not contribute measurably to substrate classification, indicating that Dis3l2 engagement and threading depend primarily on 3′-proximal sequence composition and uridine dosage rather than terminal nucleotide identity.

Together, these results reveal that Dis3l2 substrate selection follows a compositional code in which the number and position of 3′-proximal uridines tune the trade-off between binding-driven engagement and efficient threading, with optimal decay achieved at approximately four uridines.

## Discussion

Post-transcriptional uridylation has long been linked to RNA turnover, yet the mechanistic logic by which uridylation commits RNAs to degradation has remained unclear. Our study establishes that uridylation-mediated RNA decay is governed by a substrate-encoded logic, in which the kinetic properties of tail synthesis define RNA 3′-end states that are selectively decoded by the exoribonuclease, rather than by uridylation acting as a generic decay signal.

A central insight from this work is that distributive oligouridylation by the TUTase Tailor represents a functionally optimized mode of activity rather than an inefficient precursor to polyuridylation. Tailor predominantly generates short oligo(U) tracts that persist as discrete intermediates, aligning with the logic of RNA surveillance in which tailing produces decay-competent end states rather than stable extensions. This behavior contrasts with canonical poly(A) polymerases, which synthesize long tails rapidly and processively *in vivo* when associated with stimulatory cofactors^42,43^ and instead parallels quality-control pathways such as TRAMP-mediated polyadenylation, where tail addition promotes RNA exosome recruitment and decay rather than stabilization^30^.

Our data reveal two separable but complementary, enzyme-intrinsic mechanisms that together constrain U-tail length during Tailor-mediated uridylation. First, Tailor dynamically modulates its kinetics within the oligouridylation regime by sensing the emerging U-tail: as short oligo(U) tracts accumulate, elongation progressively slows, biasing the reaction toward discrete intermediates rather than uniform extension. Such product-dependent modulation is well precedented among terminal transferases. Structural studies of trypanosomal TUTases showed that RNA primer geometry, base stacking, and transient 3′-end disengagement directly influence repeated nucleotide addition^44,45^, while work on the Cid1 family demonstrated that nucleotide selectivity is biased rather than absolute and can be tuned by subtle conformational features of the enzyme–RNA complex^46^. Second, Tailor’s relaxed nucleotide selectivity imposes an additional and mechanistically distinct constraint that prevents acquisition of a sustained processive state under physiological mixed-NTP conditions. Sporadic non-uridine incorporation events are strongly enriched at tail termini and are rarely followed by further extension, effectively acting as chain-attenuating events. Kinetic modeling further shows that suppression of tail elongation cannot be explained by misincorporation alone but also reflects non-productive co-substrate competition—particularly by excess ATP—that slows catalysis and disfavors prolonged processivity. Consistent with this principle, studies of the fission yeast TUTase Cid1 showed that nucleotide identity modulates the balance between distributive and extended tailing through changes in enzyme–RNA affinity and conformational control, rather than through misincorporation per se^47^. Together, these two intrinsic layers—tail-length–dependent kinetic tuning within oligouridylation and co-substrate–mediated suppression of sustained processivity—explain how Tailor robustly generates short, decay-competent oligo(U) intermediates while intrinsically disfavoring entry into a polyuridylation mode.

On the RNA decay side, our quantitative analysis defines how Dis3l2 selectively engages Tailor-primed substrates. Dis3L2 degrades structured RNAs without ATP-dependent helicases, distinguishing it from the RNA exosome and functionally placing it closer to bacterial RNase II–family enzymes that couple binding, unwinding, and degradation within a single polypeptide^29,48,49^. Structural studies revealed that Dis3L2 contains an extended oligo(U)-binding path and undergoes large conformational rearrangements upon engagement of structured RNA, exposing a trihelix linker that promotes strand separation and helicase-independent degradation^26,27^. Our data provide a quantitative framework for how uridylated substrates are admitted into this pathway despite the apparent disparity between short 3′ overhangs and an extended RNA-binding channel. Massively parallel binding and decay measurements reveal a sharp gating step for Dis3l2: productive decay requires a minimal accessible 3′ overhang of approximately four nucleotides, corresponding to a discrete transition from binding to efficient threading and degradation. This threshold is dictated by 3′ proximal uridine content, local accessibility, and threading competence. Thus, Dis3l2 integrates multiple substrate features before committing RNAs to processive decay. These findings explain how short oligo(U) tails can satisfy the requirements of Dis3l2’s extended RNA-binding path. Initial threading of a minimal single-stranded overhang appears sufficient to pass the entry gate, while distributed uridine contacts stabilize the threaded state as the RNA progresses toward the catalytic site. Occupation of the full uridine-binding path is likely achieved progressively during translocation, consistent with the conformational plasticity observed for Dis3L2^27^. Notably, the tolerance of non-uridine residues at the extreme 3′-end further ensures that Dis3l2 can robustly account for the co-substrate promiscuity of Tailor-mediated uridylation.

Collectively, our findings support a substrate-centric model for uridylation-mediated RNA surveillance. In *Drosophila*, Tailor and Dis3l2 have been reported to physically interact, yet our results indicate that decay specificity can be explained by RNA-encoded features without invoking obligate enzyme–enzyme coupling. This logic readily accommodates mammalian systems, where TUT4/7 and Dis3L2 cooperate in cytoplasmic RNA surveillance without clear evidence for stable complex formation^21-23^.

More broadly, our findings suggest that RNA quality control can be enforced through transient, kinetically encoded RNA end states rather than stable, categorical modification marks. In this framework, decay competence emerges only within a narrow temporal window defined by tail length, composition, and accessibility, and requires productive decoding by the exoribonuclease.

### Limitations of the study

This study dissects the kinetic principles underlying uridylation-mediated RNA decay using reconstituted biochemical systems and massively parallel *in vitro* assays. While this approach enables quantitative resolution of tailing and decay reactions, it necessarily abstracts enzymatic activities from the full cellular context. *In vivo*, additional regulatory factors, RNA-binding proteins, and subcellular compartmentalization may modulate Tailor and Dis3l2 activities, influence substrate availability, or reshape kinetic parameters defined here. How such factors tune uridylation and decay under physiological or stress conditions remains to be determined.

Our analyses focus on defined RNA substrates with controlled sequence and structural features, allowing systematic interrogation of kinetic dependencies but not capturing the full diversity of endogenous RNA architectures or RNA–protein assemblies. Moreover, although Tailor and Dis3l2 physically interact in *Drosophila*, our study emphasizes functional coupling through substrate features rather than direct enzyme–enzyme interactions; whether and how physical association contributes to regulation or efficiency *in vivo* was not addressed.

Finally, while the quantitative framework developed here reveals general principles of uridylation-coupled decay, its applicability to other uridylation pathways, RNA classes, or organisms will require further experimental validation.

## Resource Availability

### Lead Contact

Requests for further information and resources should be directed to and will be fulfilled by the lead contact, Stefan L. Ameres

### Materials Availability

Reagents generated in this study will be made available on request.

### Data and Code Availability

Original gel images are uploaded to Zenodo and can be accessed via the following link: https://doi.org/10.5281/zenodo.18223078.

Raw sequencing data have been deposited at ENA (accession: PRJEB102466) and are publicly available as of the date of publication.

Processed sequencing data are uploaded to Zenodo and can be accessed via the following link: https://doi.org/10.5281/zenodo.17925051.

All original code has been deposited at GitHub at github.com/SenseAI/uridylation and is publicly available as of the date of publication.

Any additional information required to reanalyze the data reported in this paper is available from the lead contact upon request.

## Acknowledgments

We thank all members of the Ameres laboratory for helpful discussions; Ulrich Hohmann and the Protech facility for help with the purification of recombinant Tailor from *Sf9* insect cells; and the IMP/IMBA/GMI and Max Perutz Labs core facilities for the excellent support. HTP sequencing was performed at the VBCF NGS Unit (www.vbcf.ac.at). This work was supported by an EMBO Long Term Fellowship (ALTF 121-2019) and an FWF ESPRIT (10.55776/ESP591) to A.S. Research in the S.L.A. laboratory is supported by the European Union (European Research Council; RiboTrace, CoG-866166), the Austrian Science Fund (FWF; 10.55776/F80 and 10.55776/DOC177) and the Vienna Science and Technology Fund (WWTF; LS23-053).

## Author Contributions

A.S. and S.L.A. conceived the project and designed the experiments. A.S. conducted all experiments. F.B. and M.J. purified recombinant Dis3l2^CM^ (Δ51-200). A.S., D.M. and T.R.B. performed bioinformatics analysis of high-throughput sequencing datasets. D.M., B.M.J. and A.A. performed kinetic modeling. N.P. performed machine learning analyses. A.S. and S.L.A. wrote the manuscript with input from all authors.

## STAR Methods

### Experimental Model and Subject Details

#### S2 cells

FLAG-tagged *Drosophila melanogaster* Dis3l2 was expressed in S2 cells and immunopurified as described previously^20^.

#### Sf9 insect cells

Recombinant *Drosophila melanogaster* Tailor and Dis3l2^CM^ proteins were expressed in Sf9 cells using the Bac-to-Bac™ baculovirus expression system (Invitrogen).

#### Drosophila embryo lysates

Endogenous Tailor protein levels were assessed in *Drosophila melanogaster* 0–2 h embryo lysates (Figure S1A,B).

## Method Details

### Quantification of Tailor Protein Levels in Embryos

#### Western blotting

Proteins from 0–2 h embryo lysates were resolved by SDS–PAGE and transferred to nitrocellulose. Membranes were blocked in PBS containing 5% (w/v) fat-free milk and 0.3% Tween-20, then incubated with anti-Tailor monoclonal antibody (1:500 dilution), which detects both recombinant and endogenous Tailor species^20^. Signals were detected using the ECL Western Blotting Detection System (GE Healthcare) according to the manufacturer’s instructions.

#### Estimation of intracellular Tailor concentration

Tailor abundance in 0–2 h embryo lysates was quantified by western blot using the endogenous anti-Tailor antibody and a recombinant Tailor standard curve. Band intensity from 60 µg embryo lysate was interpolated on the standard curve to yield 5.27 ng Tailor in the sample. Assuming that 3.2% of embryo weight is protein and that a total embryo weight of 1,875 µg corresponds to a volume of 1,875 nL, intracellular Tailor concentration was estimated to be ∼40 nM^50^.

### Protein Expression and Purification

#### Recombinant Tailor ([His]_6_–MBP–Tailor)

Cloning, baculovirus generation, and production of infected Sf9 cultures were performed by the Protein Technologies (ProTech) Facility at Vienna BioCenter Core Facilities (VBCF). The Tailor coding sequence was amplified from plasmid template and subcloned into pGB in frame with an N-terminal HRV 3C–cleavable (His)_6_–MBP affinity tag^51^. Pellets from 1 L culture were resuspended in 100 mL lysis buffer (25 mM HEPES pH 7.5, 500 mM NaCl, 0.1% Tween-20, 2 mM DTT) supplemented with protease inhibitor cocktail (Roche) and 20 mM imidazole, lysed by sonication, and clarified by ultracentrifugation at 4 °C. Cleared lysate was loaded onto an HisTrap column (Cytiva/GE Healthcare Life Sciences) equilibrated in lysis buffer at 1 mL/min. Eluted protein was further purified on a HiTrap SP HP column (Cytiva) equilibrated in 25 mM HEPES pH 7.5, 5% glycerol, 2 mM DTT, eluting over ∼125 mL with a 0–1 M NaCl gradient. Fractions were assessed by SDS–PAGE, concentrated (VivaSpin 20, 30 kDa MWCO), and subjected to size exclusion chromatography (Superdex 200 pg 16/600, Cytiva) equilibrated in SEC buffer (30 mM HEPES pH 7.4 [KOH], 100 mM KOAc, 2 mM Mg(OAc)_2_, 5% glycerol, 1 mM TCEP). Peak fractions were analyzed by SDS–PAGE, concentrated, and stored at 4 °C for short term or flash-frozen in liquid nitrogen and stored at −80 °C for long term.

#### FLAG–Dis3l2 (S2 cell immunopurification)

FLAG-tagged Dis3l2 was immunopurified from S2 cells as previously described^20^.

#### Dis3l2^CM^ (Δ51–200, E569A; catalytically inactive)

A catalytically inactive *Drosophila melanogaster* Dis3l2 variant (E569A) lacking residues 51–200 (Dis3l2^CM^, Δ51–200) was expressed in Sf9 cells using Bac-to-Bac™ as a fusion bearing N-terminal TwinStrep and SUMOstar tags separated by an HRV 3C protease cleavage site. Sf9 cells were harvested 64 h post-infection, washed with PBS, and resuspended in lysis buffer (25 mM Tris pH 8.0, 500 mM NaCl, 0.4% Triton X-100, 5 mM DTT) supplemented with 1 µg/mL DNase I and cOmplete™ protease inhibitor cocktail (Roche 06538282001; 50 mL per 1 L culture). Cells were lysed by sonication and clarified by centrifugation. Lysate was incubated with Strep-Tactin beads (IBA 2-1000-005) for 1 h at 4 °C. Beads were washed with high-salt buffer (25 mM Tris pH 8.0, 2 M NaCl, 5 mM DTT, 0.1% Tween-20), followed by a wash in buffer containing 200 mM NaCl. Bound protein was eluted in 20 mM Tris pH 8.0, 200 mM NaCl, 5 mM DTT, 5 mM desthiobiotin and cleaved overnight with HRV 3C protease at 4 °C. Following cleavage, the NaCl concentration was reduced to 100 mM using 25 mM Tris pH 8.0, 5 mM DTT. Cleaved protein was purified by cation exchange chromatography on a HiTrap Heparin column (Cytiva 17-0407-03) using a 0.1–2 M NaCl gradient in 25 mM Tris pH 8.0, 5 mM DTT. Final purification was performed by size exclusion chromatography (Superdex 200 Increase 10/300, Cytiva 28990944) in 25 mM HEPES pH 8.0, 500 mM KCl, 5 mM DTT, 2 mM MgCl_2_. Fractions were concentrated, flash-frozen, and stored at −80 °C.

### RNA Substrate Preparation

#### Synthetic RNAs and radiolabeling

All synthetic RNAs were purchased from Horizon (sequences in Table 2). For radiolabeling, 30 pmol RNA was 5′-end labeled with γ-^32^P-ATP (6,000 Ci/mmol; Hartmann Analytic) using T4 polynucleotide kinase (PNK; NEB) and purified using G25 spin columns (GE Healthcare). Labeled RNAs were gel-purified on 10% or 15% denaturing PAGE.

### *In Vitro* RNA Uridylation Assays

#### Denaturing PAGE tailing assays

Tailing reactions were performed at 25 °C and contained recombinant (His)_6_–MBP–Tailor (30 nM) with 10 nM 5′-^32^P-labeled RNA substrate (single-stranded substrates with either 4 random nucleotides at the 3′-end, 6N, or as indicated in figure legends), including 0.5 mM UTP^52^. To set up the reaction, protein and RNA were mixed with 3 μl 40 x reaction mix (20 mM DTT, 1.67 mM UTP, 160 mM KOAc, 11.2 mM Mg(OAc)_2_) and 4 μl 1 x Lysis buffer (30 mM HEPES-KOH pH 7.4, 100 mM KOAc, 2 mM Mg(OAc)_2_, 5 mM DTT) in a total reaction volume of 10 μl. For Figure 2 experiments, reactions were additionally performed under (i) equal NTP conditions (0.5 mM each NTP) or (ii) physiological NTP conditions (0.5 mM UTP, 0.5 mM GTP, 0.3 mM CTP, 3 mM ATP). In both settings, Mg(OAc)_2_ was adjusted to account for NTP concentrations. Reactions were initiated by Tailor addition. Aliquots were removed at indicated times, quenched with formamide dye, resolved by denaturing PAGE at single-nucleotide resolution, dried, visualized on a Storm PhosphorImager (GE Healthcare), and quantified with ImageQuant TL 8.1 (GE Healthcare).

#### Scavenger assays to assess stepwise processivity

To determine stepwise uridylation processivity, reactions were assembled as above using 10 nM 5′-^32^P-labeled single-stranded RNA and 30 nM Tailor. At defined times after reaction initiation (t_1_), scavenger RNA was added to 10 µM to test rebinding of Tailor to substrate. Following scavenger addition, aliquots were removed at defined times and processed on 15% denaturing PAGE at single-nucleotide resolution. Gels were dried, imaged on a Typhoon, and quantified using ImageQuant TL 8.1.

### High-Throughput Biochemical Assays

#### RNA library construction

Libraries for high-throughput biochemical tailing and RNA decay assays were generated as previously described^20^. Briefly, RNA was recovered from the *in vitro* RNA uridylation assays by phenol-chloroform extraction and ligated to 3′ adapter containing 4 random nucleotides at the ligation interface to minimize ligation bias^53^. SuperScript III Reverse Transcriptase (Invitrogen) was used for reverse transcription, and the cDNA samples were PCR amplified using KAPA Reat-Time Library Amplification Kit (Roche). Amplified cDNA was purified on 2% agarose gels, followed by library quality control and high-throughput sequencing performed by the VBCF Next Generation Sequencing facility.

#### Tail composition analysis in HT tailing experiments with all NTPs

For HT tailing experiments with all NTPs (Figure 2), tail RPM values were summed across biological replicates (UTP-only: n=3; equal NTPs: n=4; physiological NTPs: n=3). For Figure 2E, average tail length was simulated using a chain-termination model. Uridylation rates (*k*_1_–*k*_10_) averaged over all substrates were used for uridylation steps. Non-U incorporation rates were set to 0 (UTP-only), 3.8% of the corresponding uridylation rates (equal NTPs), or 5.2% of the corresponding uridylation rates (physiological NTPs), based on observed non-U incorporation at 0.833 min. For Figure S2H,I, tails were aligned by their 5′- or 3′-ends, and per-position non-U fractions were determined from summed tail RPMs across all tail lengths and replicates (equal NTPs: n=4; physiological NTPs: n=3).

### *In Vitro* RNA Decay Assays

#### Dis3l2 decay reactions

RNA decay reactions contained 10 nM FLAG–Dis3l2 and 10 nM 5′-^32^P-labeled RNA substrate (single-stranded RNA with 6 random nucleotides at the 3′-end) at 25 °C, including 0.5 mM UTP^52^. To set up the reaction, protein and RNA were mixed with 3 μl 40 x reaction mix (20 mM DTT, 1.67 mM UTP, 160 mM KOAc, 11.2 mM Mg(OAc)_2_) and 4 μl 1 x Lysis buffer (30 mM HEPES-KOH pH 7.4, 100 mM KOAc, 2 mM Mg(OAc)_2_, 5 mM DTT) in a total reaction volume of 10 μl.

### RNA Bind-n-Seq

#### Double-filter equilibrium binding assay

RNA binding assays were performed essentially as described^54^. Double-filter binding was used to measure equilibrium binding of Dis3l2_CM_ (Δ51–200) to a single-stranded RNA substrate containing 6 random nucleotides at the 3′ end. To prevent degradation, catalytically inactive Dis3l2_CM_ was used throughout. Binding reactions (10 µL) were assembled in 30 mM HEPES-KOH pH 7.9, 120 mM KOAc, 3.5 mM Mg(OAc)_2_, 2 mM DTT, 1 U/µL SUPERase•In (Thermo Fisher), and 2.5 µg/µL BSA, containing 10 nM 5′-^32^P-labeled RNA and increasing concentrations of recombinant Dis3l2^CM^ (0.07–4,000 nM). A no-protein control was included to account for background membrane binding and was sequenced as input. Reactions were incubated for 2 h at 25 °C to reach equilibrium. Protein–RNA complexes were retained on Protran nitrocellulose membranes (Whatman/GE Healthcare), and unbound RNA was captured on Hybond-N+ membranes (Cytiva) using a Bio-Dot apparatus (Bio-Rad). After vacuum application, membranes were washed three times with 100 µL cold wash buffer (30 mM HEPES-KOH pH 7.9, 120 mM KOAc, 3.5 mM Mg(OAc)_2_, 5 mM DTT), separated, and air-dried. Bound RNA was visualized by phosphorimaging. Nitrocellulose pieces containing bound RNA were excised and digested with Proteinase K (1 µg/µL; Thermo Fisher) in 100 mM Tris-HCl pH 7.5, 10 mM EDTA, 150 mM NaCl, 1% (w/v) SDS for 1 h at 45 °C with shaking (300 rpm). A synthetic RNA spike-in (5′-ACACUCUUUCCCUACACGACGCUCUUCCGAUCU-3′; Horizon) was added at this step (expected 1:100 ratio) to enable quantitative comparison across binding reactions; the spike-in was gel-purified on 10% denaturing PAGE. RNA was recovered by phenol–chloroform extraction and ethanol precipitation, then reverse transcribed, amplified, and sequenced as described above.

### Quantification and Statistical Analysis

#### Gel quantification

Gel images were quantified using ImageQuant TL 8.1 (GE Healthcare).

#### RNA decay rate estimation (*k*_deg_)

For gel-based decay experiments, the fraction of remaining RNA at each timepoint was determined by band quantification. For HT decay datasets, RPM values for each RNA species were multiplied by the remaining fraction at each timepoint, normalized to the 0 min timepoint, and fit to a single-exponential decay model using nls() in R (v4.4.2; 2024-10-31 ucrt):

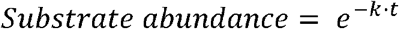

with *k* = 1 as initial guess. RNA species yielding negative degradation rate constants (n=62) were excluded from downstream analyses.

#### RNA Bind-n-Seq analysis and *K*_d_ determination

Spike-in counts were normalized by dividing by total read counts. Reads for each substrate were normalized to spike-in by dividing substrate RPM by normalized spike-in counts. These values were then divided by the corresponding normalized RPM in the input (no-protein) sample to obtain relative abundances for each substrate. Because higher membrane retention was observed in the absence of protein than at the lowest protein concentration (0.07 nM), background binding was not subtracted. Relative abundances were averaged across replicates and fit to:

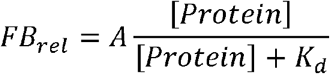

using nls.lm() from the minpack.lm package package^55^ in R (v4.4.2; 2024-10-31 ucrt)^56-58^. Initial conditions were set to the maximum observed relative fraction bound for A and the half-maximal protein concentration for *K*_d_. Parameter A was unconstrained; *K*_d_ was constrained to be non-negative.

#### Definition of Productive Score (PS) and Threading Score (TS)

To integrate RNA binding affinity and decay efficiency into composite metrics describing Dis3l2 substrate handling, decay rate constants (*k*_deg_) and dissociation constants (*K*_d_) were first standardized across the full substrate pool using robust Z-score normalization. Robust Z-scores were calculated by subtracting the median and dividing by the median absolute deviation (MAD) for each parameter independently. The productive score (PS) was defined as:

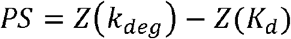

such that high PS values identify substrates that combine strong binding (low *K*_d_) with efficient degradation (high *k*_deg_).

To capture decay efficiency relative to binding strength and thus report on the ability of bound substrates to support productive RNA translocation into the catalytic site, we additionally defined a threading score (TS) as:

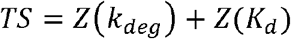

In this formulation, high TS values indicate substrates that decay rapidly despite weaker binding, consistent with efficient threading and translocation, whereas low TS values indicate substrates whose decay is limited relative to their binding affinity.

Together, PS and TS decouple substrate binding, productive engagement, and translocation efficiency and were used throughout Figure 4 and Figure S4 analyses.

#### Productive score positional contribution and cooperativity analysis

To quantify how individual 3′-nucleotide positions and their pairwise combinations influence productive RNA decay, we analyzed randomized 6N substrates predicted to be unstructured (−EFE ≤ 1.5 kcal mol^−1^). All analyses were performed on productive score (PS) values. For Individual positional contributions at each position (P1–P6), an individual positional contribution (*ΔPS*_*i*_) was calculated as the shift in median PS for substrates containing a specific nucleotide at position *i* relative to the global median PS across all analyzed substrates. This metric captures the additive, context-independent contribution of a single specific nucleotide at a given position. Pairwise cooperativity between uridines at positions *i* and *j* was quantified using the Cooperativity of Productive Score (CPS). For each position pair, a test group comprised substrates containing uridines at both positions, while a control group comprised substrates containing a uridine at only one of the two positions, with total uridine content constrained between 1 and 5 to control for uridine dosage. The observed effect (ΔPS_obs_) was defined as the median PS difference between test and control groups. The expected additive effect was calculated as the mean of the corresponding individual positional contributions:

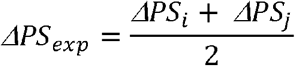

CPS was then defined as:

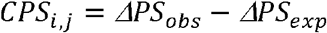

Statistical significance was assessed using a two-sided Mann–Whitney U test comparing PS distributions of test and control groups; only CPS values with p < 0.05 were retained.

#### Sequence logo generation

To visualize nucleotide preferences across the 6N region for tailing and decay, substrates in the top 10% or bottom 10% of rate constants were selected (tailing: 410/4,096; decay: 403/4,034). For Bind-n-Seq, the 410 substrates with lowest (or highest) *K*_d_ values were selected. Position probability matrices (PPMs) were computed from selected sequences. Information content (I^seq^) was calculated following Stormo *et al*. (2000) using the RPM-weighted input PPM as background^59,60^. Negative information content values (nucleotide depletion relative to input) were removed from plots. Logos were generated using ggseqlogo() (v0.26; R)^61^. For Figure S4P, top/bottom 10% Z scores (110/1,103) were used; for Figure S4S, all substrates within each quadrant (Q1–Q4) were used.

#### Machine learning

For assessing the influence of substrate sequence and secondary structure on a given target variable (*k*_1_ or productive score), we trained a deep learning model and assessed relative permutation feature importance. Briefly, we first trained a small multilayer perceptron (MLP) with *N*_*H*_ = 2 hidden linear layers, ReLU activations and 10% dropout for regularization on the following normalized input features: *b*_*x*_ (base identity of the x^th^ nucleotide in the random postfix sequence, one-hot encoded), *pN*_*x*_ (base pairing probability of the x^th^ nucleotide) and negative EFE (free energy of the thermodynamic ensemble, Z-Score normalized) as predicted by RNAfold v2.7.0^62^. The target variable was log-transformed (ln(1+x)) to reduce skewness and stabilize variance. We trained individual models for 6N datasets where appropriate with pytorch v2.5.1 and pytorch lightning v2.5.0 (https://github.com/Lightning-AI/lightning), using a mean square error (MSE) loss function to regress on the target variables^63^. The input data was randomly split into training (70%), validation (15%) and test (15%) subsets and training was conducted using the Adam optimizer with a fixed learning rate (1E-4) for a maximum of 3000 epochs (with early stopping when no validation loss reduction was observed for 250 epochs) on a MacBook Pro M3 Max with 64GB of RAM using its MPS (Metal Performance Shaders) backend. Finally, we estimated relative permutation feature importance for each model by randomly permutating the values for each input feature in the respective dataset, predicting the target variable from the permutated data and calculating average mean loss per batch (batch size: 10). We repeated this procedure 25X per input feature to measure variance in the estimated feature importances and plotted the data.

#### Statistical software

Statistical analyses were performed in Prism v10.1.1 (GraphPad), Excel v16.80 (Microsoft), or R (v4.4.2; 2024-10-31 ucrt).

### Bioinformatics Analysis

#### Processing of HT tailing/decay/binding reads

For high-throughput characterization of Tailor and Dis3l2 substrate preferences, the targeted substrate sequence excluding random nucleotides (N) was used for sequence cutting. For sequences lacking random nucleotides (spike-in), the last nucleotide was treated as a pseudo-N. Reads were trimmed using cutadapt v1.8.3^64^: 5′ anchor (^) removal followed by 3′ adapter trimming (AGATCGGAAGAGCACACGTCTGAACTCCAGTCACNNNNNN ATCTCGTATGCCGTCTTCTGCTTG) with a minimum trimmed length of 6 nt and error rate 0.1. Only trimmed sequences were processed further. fastx_toolkit v0.0.14 (http://hannonlab.cshl.edu/fastx_toolkit/) was used to remove the 3′ k-mer by length and to filter reads requiring quality >20 at each position. Substrates (5′ k-mer) and corresponding tails were defined by k-mer length and counted using a custom Perl script. Tail contamination was estimated as described^20^. For poly(U) tail analyses, tails containing non-U nucleotides were removed. For decay/binding experiments, sequences with tails and shortened substrates were removed.

### Mathematical Modeling and Implementation

#### First-order kinetic model of Tailor-mediated uridylation

Tailor-mediated uridylation was modeled as sequential addition of uridines to RNA substrates of tail length n (0 to N). The full reaction scheme comprises Tailor binding to substrate, nucleotide addition, and unbinding. Under assumptions that (i) Tailor binding is instantaneous, (ii) Tailor concentration is constant during the reaction, and (iii) UTP is in large excess relative to substrate, the system reduces to an irreversible first-order extension model:

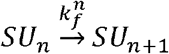

with *n* = 0,…,*N*, where *SU*_*n*_ denotes substrate with tail length *n* and 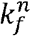 the effective first-order uridylation rate constant for step *n* ⍰ *n* +1. Sequential kinetics were represented by:

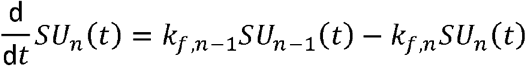

with initialization *k*_*f,-1*_=0 and *SU*_*-1*_*(t)*=0 for *n* =0. Tail lengths up to *N* =10 were modeled (11 equations, 11 initial conditions, 11 rate constants). Note, that similar first-order models have been applied to RNA adenylation and deadenylation^65,66^.

Rate constants were estimated by nonlinear least squares regression of numerical solutions to time-series data. Optimization used a constrained Nelder–Mead simplex algorithm with relative and absolute tolerances of 1×10^-7^. At each optimization step, the system was solved numerically using a 4th–5th order Runge–Kutta method with initial step size 1×10^-5^ s. Parameters were constrained to be non-negative and ≤100 s^-1^. Initial conditions were constrained to match observed starting abundances: *SU*_*0*_*(0)* =1 and *SU*_*n*_*(0)* =0 for *n* =1,…,*N*. Constraint sets differed across experimental conditions as described in the corresponding analysis sections. For each substrate, fit success required: (i) RMSE < 0.5; (ii) successful algorithm completion; (iii) ≥100 iterations; and (iv) fitted parameters not within 1×10^-5^ of their bounds.

#### Implementation and data structures

The above scheme for the estimation of the uridylation and decay rate constants was implemented using a combination of C++, TypeScript and Python. The Boost library (*Boost C++ Libraries*. URL: http://www.boost.org) was used for the numerical solution of the differential equations and the NLOpt library (*The NLopt nonlinear-optimization package*. URL: https://github.com/stevengj/nlopt) for the numerical minimization of the objective function. The code, data with tag at publication are available at the Github repository https://github.com/senseai/uridylation.

RNA-Seq data for each experiment contains the relative abundances for each substrate 2-mer (AA,AC,..,UU ) or 4-mer (AAAA,AAAC,…,UUUU), and each of their uridylated n-mers (e.g. NNNNU, NNNNUU, ) at several time points. These values are normalized to 1 by dividing the relative abundances for each of these by the total reads per million (RPM) of the n-mer at each time point. Virtual substrates were created by combining the data from groupings of n-mers. Virtual substrates were calculated based on the following patterns:

– N-letter: substrates ending with these N letters
– ALL: all substrates

Prefixes were used to denote the three different methods that were used to calculate the virtual substrate.

Underscore (_N/_NN/_NNN): To create underscore virtual substrates, the original TSV data for each time point was loaded and merged into a pandas DataFrame. The underscore virtual substrates were formed by grouping the DataFrame according to the desired substrate naming pattern and averaging the data. The result was normalized by RPM at each time point.

Dash (-N/-NN/-NNN): To create dash virtual substrates, the original TSV data for each time point was loaded and merged into a pandas DataFrame. It was then normalized by RPM at each time point. After normalization, the dash virtual substrates were formed by grouping the DataFrame according to the desired substrate naming pattern and averaging the data.

Star (*N/*NN/*NNN): Star virtual substrates are different than Underscore and Dash in that no new data is introduced, rather they refer to model solutions that are obtained by averaging the fitted parameter values across the groupings. To create the *TT substrate, fitted parameters from all substrates ending in TT are averaged. The resulting parameter values are referred to as the *TT substrate results and are used to generate data for visualization.

#### Contour plots

Contour plots were generated in R (v4.4.2; 2024-10-31 ucrt) from sequencing data or gel quantification. Simulations for Figure 3C (right panel) were performed in R using ode() from the deSolve package v1.40^67^ to solve the differential equations above with all tailing rates set to *k*_*i*_ =10 min^-1^.

## Key Resources Tables

**Table 1.**
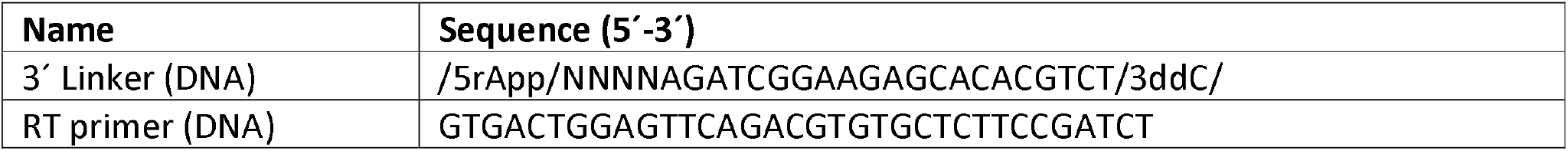
Oligonucleotides used for RNA library preparation (ddN, 2′-3′-dideoxynucleotide).

**Table 2.**
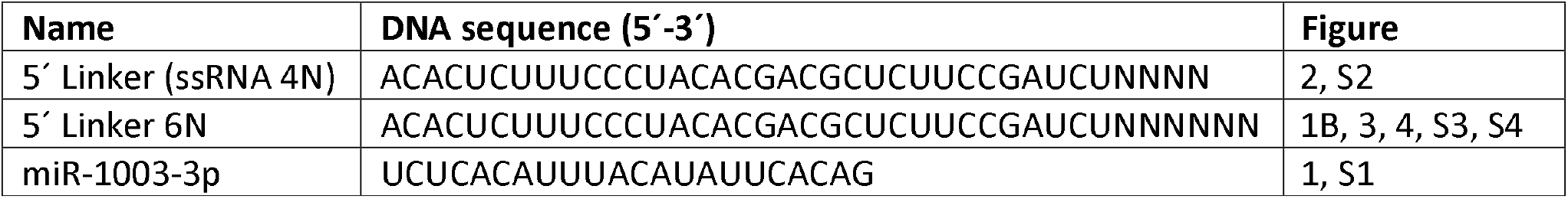
RNA oligonucleotides used for *in vitro* tailing and exoribonuclease assays.

## References

1. Houseley, J., and Tollervey, D. (2009). The many pathways of RNA degradation. Cell 136, 763–776. 10.1016/j.cell.2009.01.019.

2. Chang, H., Yeo, J., Kim, J.G., Kim, H., Lim, J., Lee, M., Kim, H.H., Ohk, J., Jeon, H.Y., Lee, H., et al. (2018). Terminal Uridylyltransferases Execute Programmed Clearance of Maternal Transcriptome in Vertebrate Embryos. Mol Cell 70, 72–82 e77. 10.1016/j.molcel.2018.03.004.

3. Curinha, A., Oliveira Braz, S., Pereira-Castro, I., Cruz, A., and Moreira, A. (2014). Implications of polyadenylation in health and disease. Nucleus 5, 508–519. 10.4161/nucl.36360.

4. Le Pen, J., Jiang, H., Di Domenico, T., Kneuss, E., Kosalka, J., Leung, C., Morgan, M., Much, C., Rudolph, K.L.M., Enright, A.J., et al. (2018). Terminal uridylyltransferases target RNA viruses as part of the innate immune system. Nat Struct Mol Biol 25, 778–786. 10.1038/s41594-018-0106-9.

5. Lim, J., Kim, D., Lee, Y.S., Ha, M., Lee, M., Yeo, J., Chang, H., Song, J., Ahn, K., and Kim, V.N. (2018). Mixed tailing by TENT4A and TENT4B shields mRNA from rapid deadenylation. Science 361, 701–704. 10.1126/science.aam5794.

6. Menezes, M.R., Balzeau, J., and Hagan, J.P. (2018). 3’ RNA Uridylation in Epitranscriptomics, Gene Regulation, and Disease. Front Mol Biosci 5, 61. 10.3389/fmolb.2018.00061.

7. Dreyfus, M., and Regnier, P. (2002). The poly(A) tail of mRNAs: bodyguard in eukaryotes, scavenger in bacteria. Cell 111, 611–613. 10.1016/s0092-8674(02)01137-6.

8. Sonenberg, N., and Hinnebusch, A.G. (2009). Regulation of translation initiation in eukaryotes: mechanisms and biological targets. Cell 136, 731–745. 10.1016/j.cell.2009.01.042.

9. Norbury, C.J. (2013). Cytoplasmic RNA: a case of the tail wagging the dog. Nat Rev Mol Cell Biol 14, 643–653. 10.1038/nrm3645.

10. Rissland, O.S., and Norbury, C.J. (2009). Decapping is preceded by 3’ uridylation in a novel pathway of bulk mRNA turnover. Nat Struct Mol Biol 16, 616–623. 10.1038/nsmb.1601.

11. Lim, J., Ha, M., Chang, H., Kwon, S.C., Simanshu, D.K., Patel, D.J., and Kim, V.N. (2014). Uridylation by TUT4 and TUT7 marks mRNA for degradation. Cell 159, 1365–1376. 10.1016/j.cell.2014.10.055.

12. Modepalli, V., and Moran, Y. (2017). Evolution of miRNA Tailing by 3’ Terminal Uridylyl Transferases in Metazoa. Genome Biol Evol 9, 1547–1560. 10.1093/gbe/evx106.

13. de Almeida, C., Scheer, H., Gobert, A., Fileccia, V., Martinelli, F., Zuber, H., and Gagliardi, D. (2018). RNA uridylation and decay in plants. Philos Trans R Soc Lond B Biol Sci 373. 10.1098/rstb.2018.0163.

14. Heo, I., Joo, C., Kim, Y.K., Ha, M., Yoon, M.J., Cho, J., Yeom, K.H., Han, J., and Kim, V.N. (2009). TUT4 in concert with Lin28 suppresses microRNA biogenesis through pre-microRNA uridylation. Cell 138, 696–708. 10.1016/j.cell.2009.08.002.

15. Heo, I., Ha, M., Lim, J., Yoon, M.J., Park, J.E., Kwon, S.C., Chang, H., and Kim, V.N. (2012). Mono-uridylation of pre-microRNA as a key step in the biogenesis of group II let-7 microRNAs. Cell 151, 521–532. 10.1016/j.cell.2012.09.022.

16. Yeom, K.H., Heo, I., Lee, J., Hohng, S., Kim, V.N., and Joo, C. (2011). Single-molecule approach to immunoprecipitated protein complexes: insights into miRNA uridylation. EMBO Rep 12, 690–696. 10.1038/embor.2011.100.

17. Reimao-Pinto, M.M., Manzenreither, R.A., Burkard, T.R., Sledz, P., Jinek, M., Mechtler, K., and Ameres, S.L. (2016). Molecular basis for cytoplasmic RNA surveillance by uridylation-triggered decay in Drosophila. EMBO J 35, 2417–2434. 10.15252/embj.201695164.

18. Lin, C.J., Wen, J., Bejarano, F., Hu, F., Bortolamiol-Becet, D., Kan, L., Sanfilippo, P., Kondo, S., and Lai, E.C. (2017). Characterization of a TUTase/RNase complex required for Drosophila gametogenesis. RNA 23, 284–296. 10.1261/rna.059527.116.

19. Bortolamiol-Becet, D., Hu, F., Jee, D., Wen, J., Okamura, K., Lin, C.J., Ameres, S.L., and Lai, E.C. (2015). Selective Suppression of the Splicing-Mediated MicroRNA Pathway by the Terminal Uridyltransferase Tailor. Mol Cell 59, 217–228. 10.1016/j.molcel.2015.05.034.

20. Reimao-Pinto, M.M., Ignatova, V., Burkard, T.R., Hung, J.H., Manzenreither, R.A., Sowemimo, I., Herzog, V.A., Reichholf, B., Farina-Lopez, S., and Ameres, S.L. (2015). Uridylation of RNA Hairpins by Tailor Confines the Emergence of MicroRNAs in Drosophila. Mol Cell 59, 203–216. 10.1016/j.molcel.2015.05.033.

21. Ustianenko, D., Pasulka, J., Feketova, Z., Bednarik, L., Zigackova, D., Fortova, A., Zavolan, M., and Vanacova, S. (2016). TUT-DIS3L2 is a mammalian surveillance pathway for aberrant structured non-coding RNAs. EMBO J 35, 2179–2191. 10.15252/embj.201694857.

22. Pirouz, M., Du, P., Munafo, M., and Gregory, R.I. (2016). Dis3l2-Mediated Decay Is a Quality Control Pathway for Noncoding RNAs. Cell Rep 16, 1861–1873. 10.1016/j.celrep.2016.07.025.

23. Labno, A., Tomecki, R., and Dziembowski, A. (2016). Cytoplasmic RNA decay pathways - Enzymes and mechanisms. Biochim Biophys Acta 1863, 3125–3147. 10.1016/j.bbamcr.2016.09.023.

24. Cheng, L., Li, F., Jiang, Y., Yu, H., Xie, C., Shi, Y., and Gong, Q. (2019). Structural insights into a unique preference for 3’ terminal guanine of mirtron in Drosophila TUTase tailor. Nucleic Acids Res 47, 495–508. 10.1093/nar/gky1116.

25. Kroupova, A., Ivascu, A., Reimao-Pinto, M.M., Ameres, S.L., and Jinek, M. (2019). Structural basis for acceptor RNA substrate selectivity of the 3’ terminal uridylyl transferase Tailor. Nucleic Acids Res 47, 1030–1042. 10.1093/nar/gky1164.

26. Faehnle, C.R., Walleshauser, J., and Joshua-Tor, L. (2014). Mechanism of Dis3l2 substrate recognition in the Lin28-let-7 pathway. Nature 514, 252–256. 10.1038/nature13553.

27. Meze, K., Axhemi, A., Thomas, D.R., Doymaz, A., and Joshua-Tor, L. (2023). A shape-shifting nuclease unravels structured RNA. Nat Struct Mol Biol 30, 339–347. 10.1038/s41594-023-00923-x.

28. Lv, H., Zhu, Y., Qiu, Y., Niu, L., Teng, M., and Li, X. (2015). Structural analysis of Dis3l2, an exosome-independent exonuclease from Schizosaccharomyces pombe. Acta Crystallogr D Biol Crystallogr 71, 1284–1294. 10.1107/S1399004715005805.

29. Matos, R.G., Garg, A., Costa, S.M., Pereira, P., Arraiano, C.M., Joshua-Tor, L., and Viegas, S.C. (2025). Structural and mechanistic insights into Dis3L2-mediated degradation of structured RNA. RNA. 10.1261/rna.080685.125.

30. Vanacova, S., Wolf, J., Martin, G., Blank, D., Dettwiler, S., Friedlein, A., Langen, H., Keith, G., and Keller, W. (2005). A new yeast poly(A) polymerase complex involved in RNA quality control. PLoS Biol 3, e189. 10.1371/journal.pbio.0030189.

31. Faehnle, C.R., Walleshauser, J., and Joshua-Tor, L. (2017). Multi-domain utilization by TUT4 and TUT7 in control of let-7 biogenesis. Nat Struct Mol Biol 24, 658–665. 10.1038/nsmb.3428.

32. Yi, G., Ye, M., Carrique, L., El-Sagheer, A., Brown, T., Norbury, C.J., Zhang, P., and Gilbert, R.J.C. (2024). Structural basis for activity switching in polymerases determining the fate of let-7 pre-miRNAs. Nat Struct Mol Biol. 10.1038/s41594-024-01357-9.

33. Han, X., Yamashita, S., and Tomita, K. (2026). Mechanistic insights into Lin28-dependent oligo-uridylylation of pre-let-7 by TUT4. Nucleic Acids Res 54. 10.1093/nar/gkaf1421.

34. Yates, L.A., Fleurdepine, S., Rissland, O.S., De Colibus, L., Harlos, K., Norbury, C.J., and Gilbert, R.J.C. (2012). Structural basis for the activity of a cytoplasmic RNA terminal uridylyl transferase. Nat Struct Mol Biol 19, 782–787. 10.1038/nsmb.2329.

35. Rissland, O.S., Mikulasova, A., and Norbury, C.J. (2007). Efficient RNA polyuridylation by noncanonical poly(A) polymerases. Mol Cell Biol 27, 3612–3624. 10.1128/MCB.02209-06.

36. Kwak, J.E., and Wickens, M. (2007). A family of poly(U) polymerases. RNA 13, 860–867. 10.1261/rna.514007.

37. Traut, T.W. (1994). Physiological concentrations of purines and pyrimidines. Mol Cell Biochem 140, 1–22. 10.1007/BF00928361.

38. Ustianenko, D., Hrossova, D., Potesil, D., Chalupnikova, K., Hrazdilova, K., Pachernik, J., Cetkovska, K., Uldrijan, S., Zdrahal, Z., and Vanacova, S. (2013). Mammalian DIS3L2 exoribonuclease targets the uridylated precursors of let-7 miRNAs. RNA 19, 1632–1638. 10.1261/rna.040055.113.

39. Chang, H.M., Triboulet, R., Thornton, J.E., and Gregory, R.I. (2013). A role for the Perlman syndrome exonuclease Dis3l2 in the Lin28-let-7 pathway. Nature 497, 244–248. 10.1038/nature12119.

40. Malecki, M., Viegas, S.C., Carneiro, T., Golik, P., Dressaire, C., Ferreira, M.G., and Arraiano, C.M. (2013). The exoribonuclease Dis3L2 defines a novel eukaryotic RNA degradation pathway. EMBO J 32, 1842–1854. 10.1038/emboj.2013.63.

41. Wu, D., Pedroza, M., Chang, J., and Dean, J. (2023). DIS3L2 ribonuclease degrades terminal-uridylated RNA to ensure oocyte maturation and female fertility. Nucleic Acids Res 51, 3078–3093. 10.1093/nar/gkad061.

42. Wahle, E. (1995). Poly(A) tail length control is caused by termination of processive synthesis. J Biol Chem 270, 2800–2808. 10.1074/jbc.270.6.2800.

43. Martin, G., Moglich, A., Keller, W., and Doublie, S. (2004). Biochemical and structural insights into substrate binding and catalytic mechanism of mammalian poly(A) polymerase. J Mol Biol 341, 911–925. 10.1016/j.jmb.2004.06.047.

44. Stagno, J., Aphasizheva, I., Aphasizhev, R., and Luecke, H. (2007). Dual role of the RNA substrate in selectivity and catalysis by terminal uridylyl transferases. Proc Natl Acad Sci U S A 104, 14634–14639. 10.1073/pnas.0704259104.

45. Stagno, J., Aphasizheva, I., Rosengarth, A., Luecke, H., and Aphasizhev, R. (2007). UTP-bound and Apo structures of a minimal RNA uridylyltransferase. J Mol Biol 366, 882–899. 10.1016/j.jmb.2006.11.065.

46. Lunde, B.M., Magler, I., and Meinhart, A. (2012). Crystal structures of the Cid1 poly (U) polymerase reveal the mechanism for UTP selectivity. Nucleic Acids Res 40, 9815–9824. 10.1093/nar/gks740.

47. Munoz-Tello, P., Gabus, C., and Thore, S. (2014). A critical switch in the enzymatic properties of the Cid1 protein deciphered from its product-bound crystal structure. Nucleic Acids Res 42, 3372–3380. 10.1093/nar/gkt1278.

48. Schneider, C., and Tollervey, D. (2014). Looking into the barrel of the RNA exosome. Nat Struct Mol Biol 21, 17–18. 10.1038/nsmb.2750.

49. Chu, L.Y., Hsieh, T.J., Golzarroshan, B., Chen, Y.P., Agrawal, S., and Yuan, H.S. (2017). Structural insights into RNA unwinding and degradation by RNase R. Nucleic Acids Res 45, 12015–12024. 10.1093/nar/gkx880.

50. Kellogg, D.R., Field, C.M., and Alberts, B.M. (1989). Identification of microtubule-associated proteins in the centrosome, spindle, and kinetochore of the early Drosophila embryo. J Cell Biol 109, 2977–2991. 10.1083/jcb.109.6.2977.

51. Neuhold, J., Radakovics, K., Lehner, A., Weissmann, F., Garcia, M.Q., Romero, M.C., Berrow, N.S., and Stolt-Bergner, P. (2020). GoldenBac: a simple, highly efficient, and widely applicable system for construction of multi-gene expression vectors for use with the baculovirus expression vector system. BMC Biotechnol 20, 26. 10.1186/s12896-020-00616-z.

52. Ameres, S.L., Horwich, M.D., Hung, J.H., Xu, J., Ghildiyal, M., Weng, Z., and Zamore, P.D. (2010). Target RNA-directed trimming and tailing of small silencing RNAs. Science 328, 1534–1539. 10.1126/science.1187058.

53. Jayaprakash, A.D., Jabado, O., Brown, B.D., and Sachidanandam, R. (2011). Identification and remediation of biases in the activity of RNA ligases in small-RNA deep sequencing. Nucleic Acids Res 39, e141. 10.1093/nar/gkr693.

54. Wee, L.M., Flores-Jasso, C.F., Salomon, W.E., and Zamore, P.D. (2012). Argonaute divides its RNA guide into domains with distinct functions and RNA-binding properties. Cell 151, 1055–1067. 10.1016/j.cell.2012.10.036.

55. Elzhov T.V. M.K.M.,, Spiess A., Bolker B. (2023). minpack.lm: R Interface to the Levenberg-Marquardt Nonlinear Least-Squares Algorithm Found in MINPACK, Plus Support for Bounds.

56. Wickham H. A.M.,, Bryan J., Chang W., McGowan L.D., François R., Grolemund G., Hayes A., Henry L., Hester J., Kuhn M., Pedersen T.L., Miller E., Bache S.M., Müller K., Ooms J., Robinson D., Seidel D.P., Spinu, and V., T.K., Vaughan D., Wilke C., Woo K., Yutani H. (2019). Welcome to the tidyverse. Journal of Open Source Software 4, 1686.

57. Wickham H. F.R.,, Henry L., Müller K., Vaughan D. (2023). dplyr: A Grammar of Data Manipulation.

58. Team, R.C. (2024). R: A Language and Environment for Statistical Computing. Austria: R Foundation for Statistical Computing.

59. Stormo, G.D. (2000). DNA binding sites: representation and discovery. Bioinformatics 16, 16–23. 10.1093/bioinformatics/16.1.16.

60. D’Haeseleer, P. (2006). What are DNA sequence motifs? Nat Biotechnol 24, 423–425. 10.1038/nbt0406-423.

61. Wagih, O. (2017). ggseqlogo: a versatile R package for drawing sequence logos. Bioinformatics 33, 3645–3647. 10.1093/bioinformatics/btx469.

62. Lorenz, R., Bernhart, S.H., Honer Zu Siederdissen, C., Tafer, H., Flamm, C., Stadler, P.F., and Hofacker, I.L. (2011). ViennaRNA Package 2.0. Algorithms Mol Biol 6, 26. 10.1186/1748-7188-6-26.

63. Ansel, J. (2024). PyTorch 2: Faster Machine Learning Through Dynamic Python Bytecode Transformation and Graph Compilation. ASPLOS’24 2, 929–947.

64. Martin, M. (2011). Cutadapt removes adapter sequences from high-throughput sequencing reads. EMBnet.journal 17, 10–12.

65. Jia, H., Wang, X., Liu, F., Guenther, U.P., Srinivasan, S., Anderson, J.T., and Jankowsky, E. (2011). The RNA helicase Mtr4p modulates polyadenylation in the TRAMP complex. Cell 145, 890–901. 10.1016/j.cell.2011.05.010.

66. Lee, Y.S., Levdansky, Y., Jung, Y., Kim, V.N., and Valkov, E. (2024). Deadenylation kinetics of mixed poly(A) tails at single-nucleotide resolution. Nat Struct Mol Biol. 10.1038/s41594-023-01187-1.

67. Soetaert K P.T.,, Setzer R.W. (2010). Solving Differential Equations in R: Package deSolve. Journal of Statistical Software, 33, 1–25.

