## Supplemental Information for "Kinetic logic of uridylation-mediated RNA decay"

**Content:**

SI Figures S1-S4

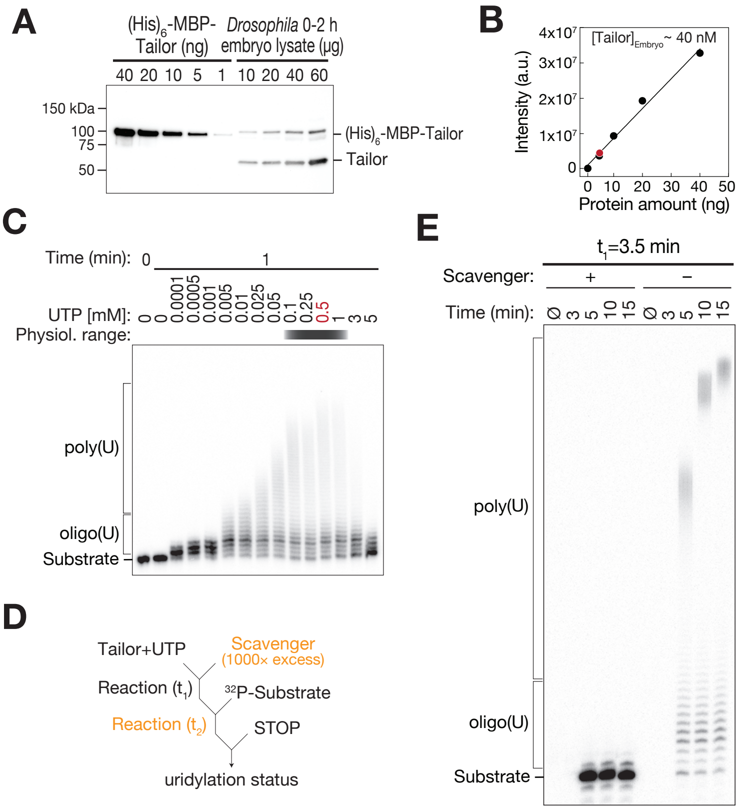


**Figure S1.** ***In vitro*** **Tailor assays approximate physiological enzyme and nucleotide conditions.** Related to Figure1.

**(A)** Immunoblot analysis of Tailor abundance in 0–2 h *Drosophila melanogaster* embryo lysates alongside recombinant $(His)_6$–MBP–Tailor. The same Tailor antibody detects endogenous and recombinant protein.

**(B)** Quantification of Tailor concentration in embryo lysates based on a recombinant Tailor standard curve. The dashed line indicates the signal corresponding to 60 µg embryo lysate, yielding an estimated intracellular Tailor concentration of ~40 nM.

**(C)** UTP titration endpoint assay (0.0001–5 mM UTP) used to define the nucleotide concentrations employed in subsequent in vitro uridylation assays. The shaded region indicates the physiological range.

**(D)** Schematic of the scavenger assay used in (E).

**(E)** Scavenger assay control demonstrating processive polyuridylation by recombinant Tailor at late time points ($t_1$ = 3.5 min), corresponding to conditions used in Figure 1G.

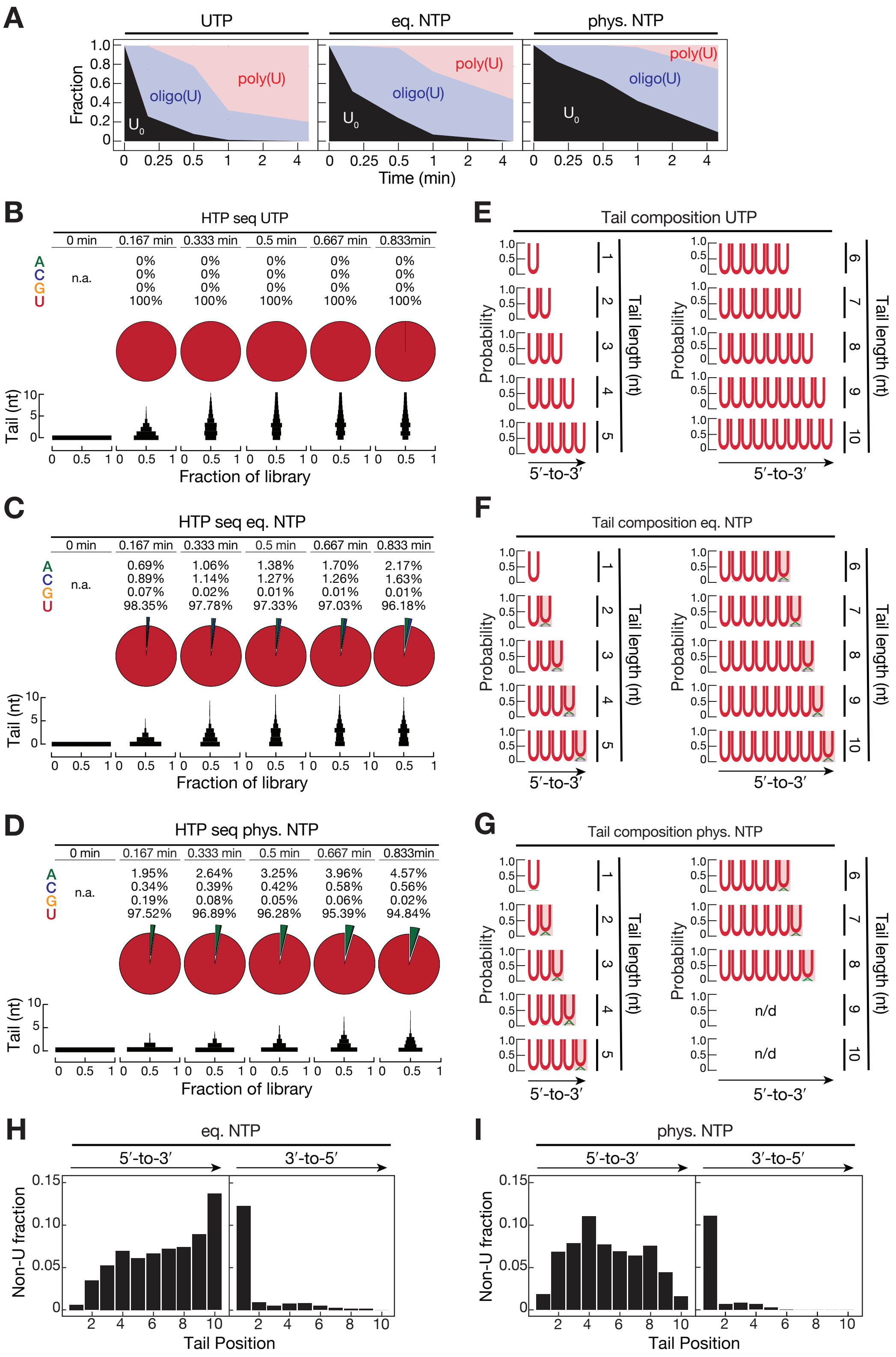


**Figure S2. Tailor displays strong uridine selectivity with terminally enriched non-uridine incorporation.** Related to Figure 2.

**(A)** Fraction of untailed substrate ($U_0$), oligo(U), and poly(U) species over time for reactions performed with UTP alone, equal NTPs, or physiological NTPs, quantified from gels shown in Figure 2A.

**(B–D)** Time-resolved nucleotide incorporation profiles for UTP-only (B), equal NTP (C), and physiological NTP (D) conditions. Pie charts show nucleotide frequencies at each time point; distributions below indicate tail length across the RNA library.
**(E–G)** Sequence logos representing tail composition (up to 10 nt) at 0.833 min for UTP-only (E), equal NTP (F), and physiological NTP (G) reactions.
**(H–I)** Positional distribution of non-uridine incorporation along RNA tails aligned from the 5′-end (left) or 3′-end (right) under equal NTP (H) and physiological NTP (I) conditions.

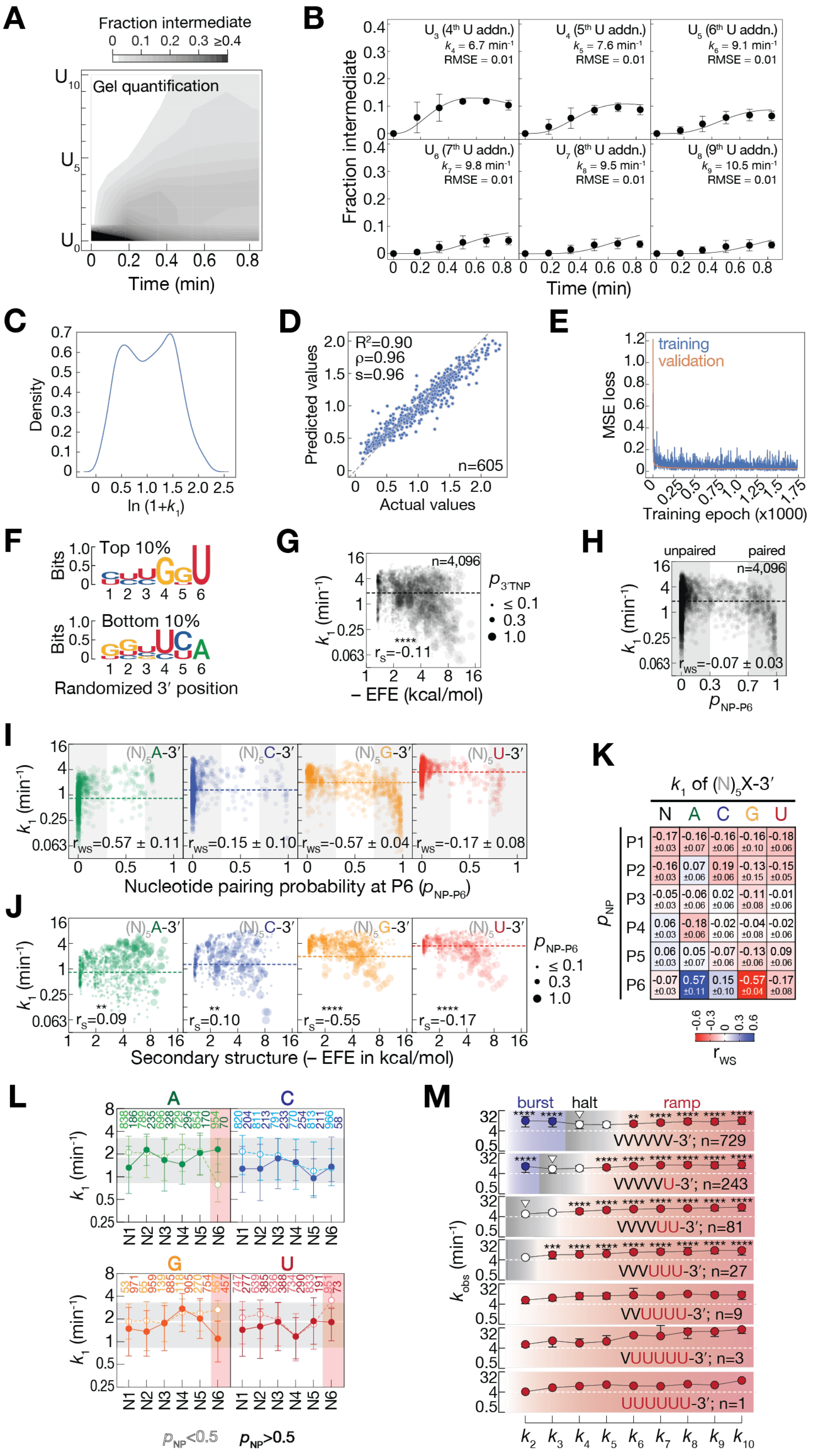


**Figure S3. Extended kinetic modeling and sequence features resolve stepwise uridylation dynamics.** Related to Figure 3.

**(A)** Contour plots showing fractions of uridylated intermediates over time based on gel quantification from three independent experiments.

**(B)** Kinetic fits for averaged normalized substrates. Mean ± SD across all 4,096 substrates are shown. Fitted rate constants ($k_{obs}$ for addition of 4–9 uridines) and RMSE values are indicated.
**(C)** Density plot of the log-transformed target variable ($ln(1+k_1)$) used for machine-learning–based prediction.
**(D)** Correlation between measured and predicted ($ln(1+k_1)$) values for the held-out test set. $R^2$, Pearson (ρ), and Spearman (s) correlation coefficients are indicated.
**(E)** Training and validation loss curves for the predictive model.
**(F)** Sequence logos representing nucleotide enrichment at the six randomized 3′ positions for the top 10% (most rapidly tailed) and bottom 10% (least rapidly tailed) substrates.
**(G)** Relationship between $k_1$ and predicted base-pairing probability of the 3′-terminal nucleotide. Dot size reflects pairing probability; dashed line indicates median $k_1$. Spearman correlation coefficient is shown.
**(H)** $k_1$ as a function of 3′-terminal nucleotide base-pairing probability for all substrates. Shaded regions denote paired and unpaired states; dashed line indicates median $k_1$. Weighted Spearman correlation coefficient is shown.
**(I)** Relationship between $k_1$ and base-pairing probability for substrates grouped by terminal nucleotide identity (A, C, G, or U). Shaded regions indicate paired and unpaired states; dashed line indicates median $k_1$. Weigthed Spearman correlation coefficient is shown.
**(J)** $k_1$ as a function of secondary structure for each terminal nucleotide identity. The size of the dots indicates the base pairing probability of the 3´terminal nucleotide (as predicted by RNAfold v2.7.0). Dashed line indicates median $k_1$. Spearman correlation coefficient is shown.
**(K)** Summary heatmap of weighted Spearman correlation coefficients between $k_1$ and base-pairing probability across terminal nucleotide identities.
**(L)** Effect of nucleotide identity across the six randomized 3′ positions on $k_1$, stratified by high (>0.5) or low (<0.5) base-pairing probability. Dashed lines indicate median $k_1$ across all substrates.
**(M)** Uridylation rates ($k_2$–$k_{10}$) for substrates lacking terminal uridines (VVVVVV) or containing increasing numbers of terminal uridines (1–6 U). White triangles denote halt conditions; adjusted p-values were calculated using Kruskal–Wallis test.

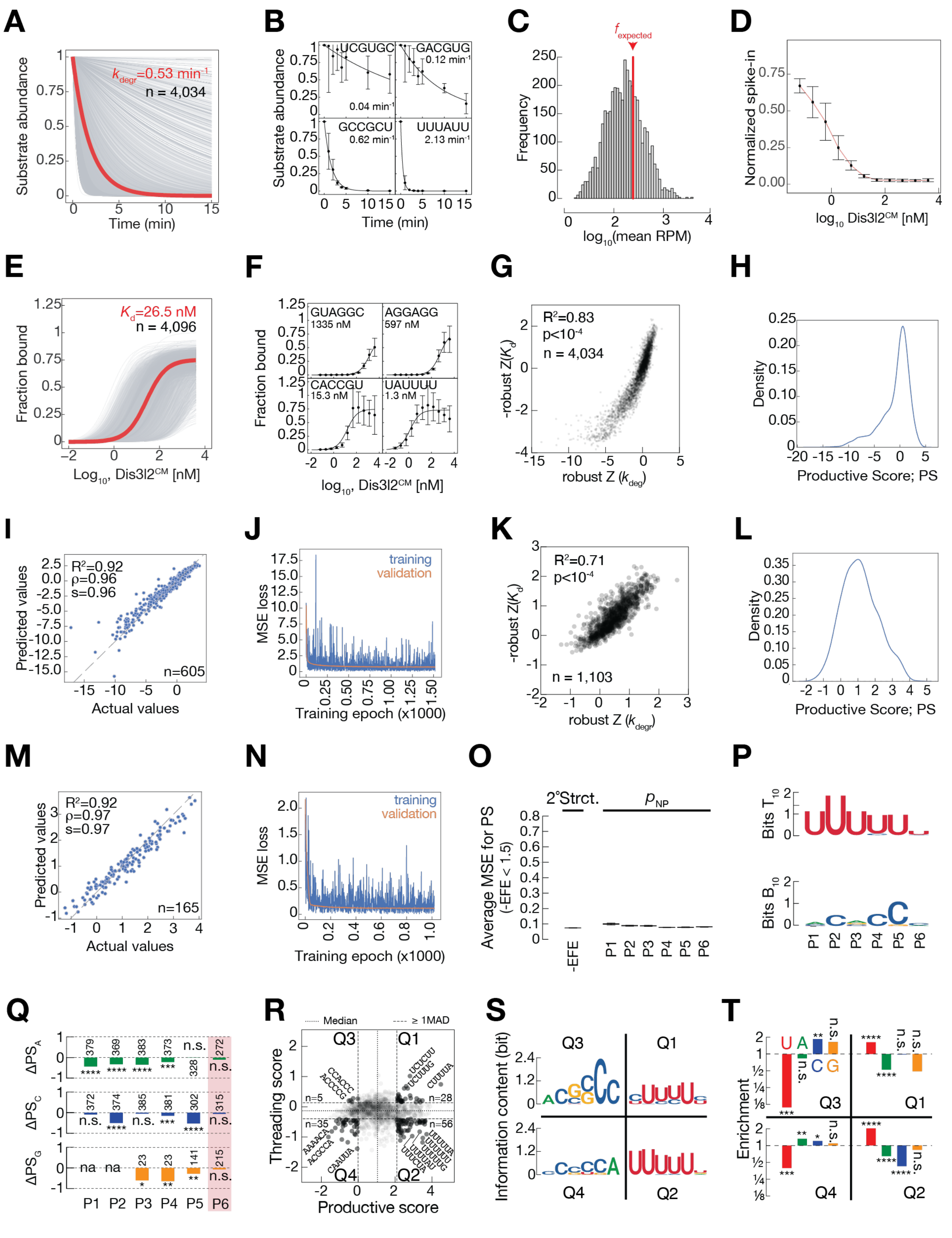

**Figure S4. Extended binding and decay analyses define substrate determinants of Dis3l2 activity.** Related to Figure 4.

**(A)** Exponential decay fits for all substrates with measurable $k_{deg}$ values; the population-averaged decay rate is shown in red.

**(B)** Representative decay profiles spanning low- to high-efficiency substrates. Data points show mean values of three replicates; error bars represent SD.

**(C)** Distribution of input substrate abundance in RNA Bind-n-Seq experiments. The RPM values for each substrate were averaged over four replicate experiments for this analysis. Statistically expected value for 4,096 equally abundant RNAs is shown with a red line.
**(D)** Normalized spike-in controls across Dis3l2$^{CM}$ concentrations.
**(E)** Global binding curves used to derive $K_d$ values for all substrates. Apparent dissociation constant of all substrates, as determined by phosphorimaging, is indicated in red.
**(F)** Representative binding curves for individual substrates. Data points show mean values of four replicates; error bars represent SD.
**(G)** Correlation between robust Z-scores of $K_d$ and $k_{deg}$ values.
**(H)** Distribution of $(ln(1 + PS))$
**(I, J)** Model performance for productive score (PS) prediction (correlation and loss curves).
**(K)** Correlation between the robust Z-scores of $K_d$ and $k_{deg}$ values for single stranded RNA substrates (-EFE$\leq$ 1.5).
**(L-N)** Corresponding analyses restricted to single-stranded substrates.
**(O)** Feature importance for single-stranded substrates (−EFE < 1.5).
**(P)** Sequence logos of the top and bottom 10% of substrates ranked by productive score (PS).
**(Q)** Position-specific contributions of adenine, cytidine and guanine to productive score (PS) in unstructured RNAs. ΔPS indicates median shifts. Significance was determined by two-sided Mann–Whitney U test; n.s., $p > 0.05$; *, $p < 0.05$; **, $p < 0.01$; ***, $p < 10^{-3}$; ****, $p < 10^{-4}$.
**(R)** Productive score (PS) versus threading score (TS) highlighting quadrant-specific outliers (±1 median absolute deviation; MAD).
**(S)** Sequence logos for substrates in each quadrant.
**(T)** Nucleotide enrichment for each quadrant (two-sided binomial tests). Significance was determined by two-sided binomial text; n.s., $p > 0.05$; *, $p < 0.05$; **, $p < 0.01$; ***, $p < 10^{-3}$; ****, $p < 10^{-4}$.
